# Mucolytic bacteria license pathobionts to acquire host-derived nutrients during dietary nutrient restriction

**DOI:** 10.1101/2022.01.17.476631

**Authors:** Kohei Sugihara, Sho Kitamoto, Prakaimuk Saraithong, Hiroko Nagao-Kitamoto, Caroline McCarthy, Matthew Hoostal, Alexandra Rosevelt, Chithra K Muraleedharan, Merritt G. Gillilland, Jin Imai, Maiko Omi, Shrinivas Bishu, John Y. Kao, Christopher J. Alteri, Nicolas Barnich, Thomas M. Schmidt, Asma Nusrat, Naohiro Inohara, Jonathan L. Golob, Nobuhiko Kamada

## Abstract

Pathobionts employ unique metabolic adaptation mechanisms to maximize their growth in disease conditions. Adherent–invasive *Escherichia coli* (AIEC), a pathobiont enriched in the gut mucosa of patients with inflammatory bowel disease (IBD), utilizes diet-derived L-serine to adapt to the inflamed gut. Therefore, the restriction of dietary L-serine starves AIEC and limits its fitness advantage. Here, we find that AIEC can overcome this nutrient limitation by switching the nutrient source from the diet to the host cells in the presence of mucolytic bacteria. During diet-derived L-serine restriction, the mucolytic symbiont *Akkermansia muciniphila* promotes the encroachment of AIEC to the epithelial niche by degrading the mucus layer. In the epithelial niche, AIEC acquires L-serine from the colonic epithelium and thus proliferates. Our work suggests that the indirect metabolic network between pathobionts and commensal symbionts enables pathobionts to overcome nutritional restriction and thrive in the gut.

## Main

Microbial metabolism plays a critical role in cooperation and competition within the microbial community^1^. Microbial metabolism rapidly responds to environmental stimulation, such as host immune activation, dietary modification, and gut inflammation, to adapt to the surrounding microenvironment^2-5^. For example, commensal symbionts reprogram the transcription of their metabolic genes in response to host immune activation and alter their metabolic functions within several hours^5^. Likewise, gut inflammation alters the luminal microenvironment, including nutrient availability and oxygen levels, which, in turn, contributes to the alteration of the gut microbial composition and function^6,7^. Metatranscriptome studies have shown that gut inflammation upregulates stress response pathways and downregulates polysaccharide utilization and fermentation in a murine model of colitis^8,9^. In addition, chronic intestinal inflammation upregulates stress response genes, including small heat shock proteins, which protect commensal *Escherichia coli* from oxidative stress^10^. These disease-specific microbial transcriptional signatures have also been observed in patients with inflammatory bowel disease (IBD)^11^. However, the impact of the transcriptional adaptation of microbes on host–microbe interaction and disease course is largely unknown.

The gut microbiota plays a fundamental role in the pathogenesis of IBD^12,13^. Potentially pathogenic members of the commensal bacteria, termed pathobionts, have been identified in IBD patients and observed to trigger or exacerbate inflammation in the gut. Adherent-invasive *Escherichia coli* (AIEC) is the most studied pathobiont associated with IBD. The prevalence of AIEC increases in the ileal and colonic mucosae of IBD patients compared to non-IBD control subjects^14^. AIEC strains harbor several virulence genes related to the ability to adhere and invade the intestinal epithelial cells (IECs) and thus are associated with the exacerbation of intestinal inflammation and fibrosis^15,16^. In IBD, AIEC may exploit unique strategies to gain a growth advantage over competing, nonpathogenic, symbiont *E. coli* strains. We have reported that AIEC reprograms metabolic gene transcription in the inflamed gut to adapt to the inflammatory microenvironment^4^. In particular, AIEC upregulates L-serine metabolism pathways that are crucial in acquiring a growth advantage over symbiont *E. coli* strains. Interestingly, as luminal L-serine is supplied by diet, the modulation of dietary L-serine can regulate intraspecific competition between AIEC and commensal *E. coli*^4^. Thus, dietary modification can be an effective strategy to treat pathobiont-driven diseases, such as IBD.

L-serine is a nonessential amino acid that supports several metabolic processes crucial for the growth and survival of mammalian and bacterial cells, especially under disease conditions^17^. For example, L-serine metabolism is markedly upregulated in cancer cells and immune cells and plays a central role in their survival and growth^18-20^. Moreover, consistent with gut bacteria, L-serine used in the proliferation of cancer cells and immune cells is also supplied by the diet, and therefore a lack of dietary L-serine can inhibit the proliferation of these cells^19,21^.

Here, we report the impact of dietary L-serine on the host–microbe interaction during gut inflammation. As the deprivation of diet-derived L-serine limits the fitness advantage of AIEC over commensal *E. coli*, we anticipated that dietary L-serine restriction would improve gut inflammation in mice colonized with conventional microbiota (specific pathogen–free [SPF] mice). However, to our surprise, diet-derived L-serine restriction exacerbated dextran sodium sulfate (DSS)–induced colitis in SPF mice. In our quest to explain this unexpected phenotype, we discovered that *Akkermansia muciniphila*, a commensal symbiont capable of degrading mucin, is expanded in colitic mice under dietary L-serine restriction. The expansion of *A. muciniphila* results in a massive erosion of the colonic mucus layer, thereby allowing AIEC to relocate close to the host epithelial cells. In the epithelial niche, AIEC can acquire L-serine from the host epithelial cells, whereby it can overcome dietary L-serine restriction and proliferate. Thus, the mucolytic bacteria, such as *A. muciniphila*, can serve as an indirect metabolic supporter for AIEC by licensing the acquisition of host-derived nutrients.

## Results

### L-serine metabolism is disturbed in the gut microbiota of IBD patients

Our previous study showed that IBD-associated AIEC uses amino acid metabolism, particularly L-serine catabolism, to adapt to the inflamed gut. Consistent with this notion, the IBD-associated AIEC strain LF82 rapidly consumed L-serine, more than other amino acids in the cultured media (**Extended Data Fig. 1**), indicating that AIEC prefers L-serine as a nutrient source. However, it remains unclear whether L-serine plays a central metabolic role in the more complex microbiota of IBD patients. To assess the microbial L-serine metabolism, we first analyzed data available in the Inflammatory Bowel Disease Multi’omics Database (IBDMBD) of the Integrative Human Microbiome Project (iHMP), which integrates metagenomic, metatranscriptomic, metaproteomic, and metabolic data on the microbiome of IBD (**Fig. 1a**). As previous studies have reported^22-24^, the abundance of Enterobacteriaceae, including *E. coli,* is significantly higher in patients with ulcerative colitis (UC) and Crohn’s disease (CD), the two most common forms of IBD, than in non-IBD controls (**Fig. 1b**). L-serine is biosynthesized from intermediates of the glycolysis pathway or from L-glycine, and it converts to pyruvate, which is a necessary substrate for gluconeogenesis and the tricarboxylic acid cycle (**Fig. 1c**, right). Metagenomic analysis showed that the abundance of phosphoglycerate dehydrogenase (PHGDH), the rate-limiting enzyme for serine biosynthesis from the glycolysis pathway, was significantly reduced in both UC and CD patients compared to non-IBD controls (**Fig. 1c**, left). Conversely, the abundance of serine dehydratase (SDH), the enzyme that catalyzes the conversion of L-serine to pyruvate, was significantly higher in the gut microbiota of UC and CD patients compared to non-IBD controls (**Fig. 1c**, left). Although metatranscriptomic profiles were more varied between individuals than metagenomic profiles, these genes were also transcriptionally changed in IBD patients (**Fig. 1d**). These results suggested that bacteria that utilize L-serine, such as AIEC, are enriched in the microbiota of IBD patients. In addition to microbial metabolism, we also confirmed the amino acid levels in the feces of patients with IBD using the iHMP metabolome data (**Fig. 1e**). Compared with non-IBD controls, essential amino acids, particularly valine and histidine, were significantly higher in patients with IBD. Importantly, nonessential amino acids, including serine, glutamate, and arginine, were significantly lower in IBD patients than in non-IBD controls. Thus, it is likely that the gut microbiota of IBD patients consumes more L-serine than the gut microbiota of non-IBD controls.

**Fig 1.**
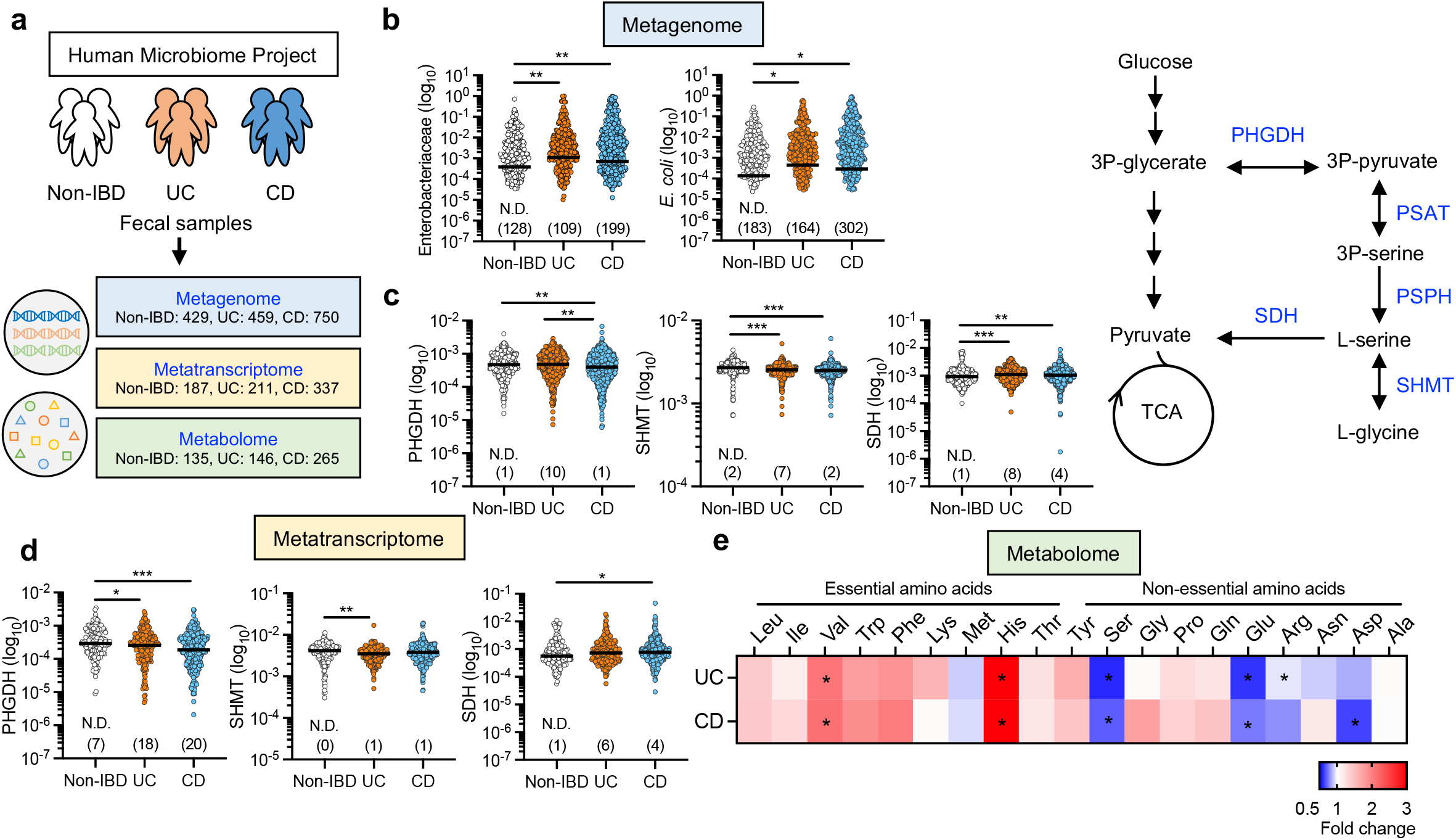
L-serine metabolism is disturbed in the gut microbiota of IBD patients. **a**, Metagenomics, metatranscriptomics, and metabolomics data were downloaded from the public resource, the second phase of the Integrative Human Microbiome Project (HMP2 or iHMP) – the Inflammatory Bowel Disease Multi’omics Database. **b**, Abundance of Enterobacteriaceae and *E. coli* in the metagenomics database. **c**, Abundance of PHGDH, SHMT, and SDH in the metagenomics database (left). Schematic of L-serine metabolism (right). **d**, Abundance of PHGDH, SHMT, and SDH in the metatranscriptomics database. **e**, Abundance of fecal amino acids in the metabolomics database. The heatmap indicates the fold change (UC or CD/non-IBD). Dots indicate individual people, with median. The numbers in parentheses indicate the number of null values. \**P* < 0.05, \*\**P* < 0.01, \*\*\**P* < 0.001 by Kruskal–Wallis test with Dunn test for multiple comparisons. PHGDH, phosphoglycerate dehydrogenase; SHMT, serine hydroxymethyltransferase; SDH, serine dehydratase. See also **Extended Data Fig. 1**.

### The deprivation of dietary L-serine exacerbates DSS-induced colitis

Given that the gut microbiota of IBD patients appeared to be enriched with L-serine utilizers, including AIEC, we hypothesized that limiting L-serine availability may suppress the growth of potential pathobionts, thereby reducing the susceptibility to colitis. As luminal L-serine levels are mainly regulated by diet-derived L-serine^4^, we next examined the impact of dietary L-serine deprivation on intestinal inflammation. To this end, specific pathogen–free (SPF) mice were fed either a defined amino acid control diet (Ctrl) or an L-serine deficient (ΔSer) diet, as previously defined^4,18^. L-glycine was removed from the ΔSer diet as L-serine and L-glycine may be interconverted^25^. Mice were treated with 1.5% DSS for 5 days to induce colitis, followed by conventional water for 2 days (**Fig. 2a**). Unexpectedly, the ΔSer diet–fed mice lost significantly more body weight and had a higher disease activity index (DAI) than the Ctrl diet–fed mice (**Fig. 2b,c**). Likewise, mice fed the ΔSer diet had a greater degree of inflammation in the colon than the mice fed the Ctrl diet (**Fig. 2d–f**). Notably, the ΔSer diet did not affect body weight, colon length, and histology in the DSS-untreated mice (**Fig. b–f**). To uncover the mechanism by which the restriction of dietary L-serine exacerbates colitis, we focused on the role of the gut microbiota. As shown in **Fig. 2g–j**, the ΔSer diet did not worsen colitis in germ-free (GF) mice. We confirmed the same phenotype in SPF mice by depleting the gut microbiota with a cocktail of broad-spectrum antibiotics (**Extended Data Fig. 2**). These results suggest that the gut microbiota is required for the exacerbation of colitis caused by the restriction of dietary L-serine.

**Fig 2.**
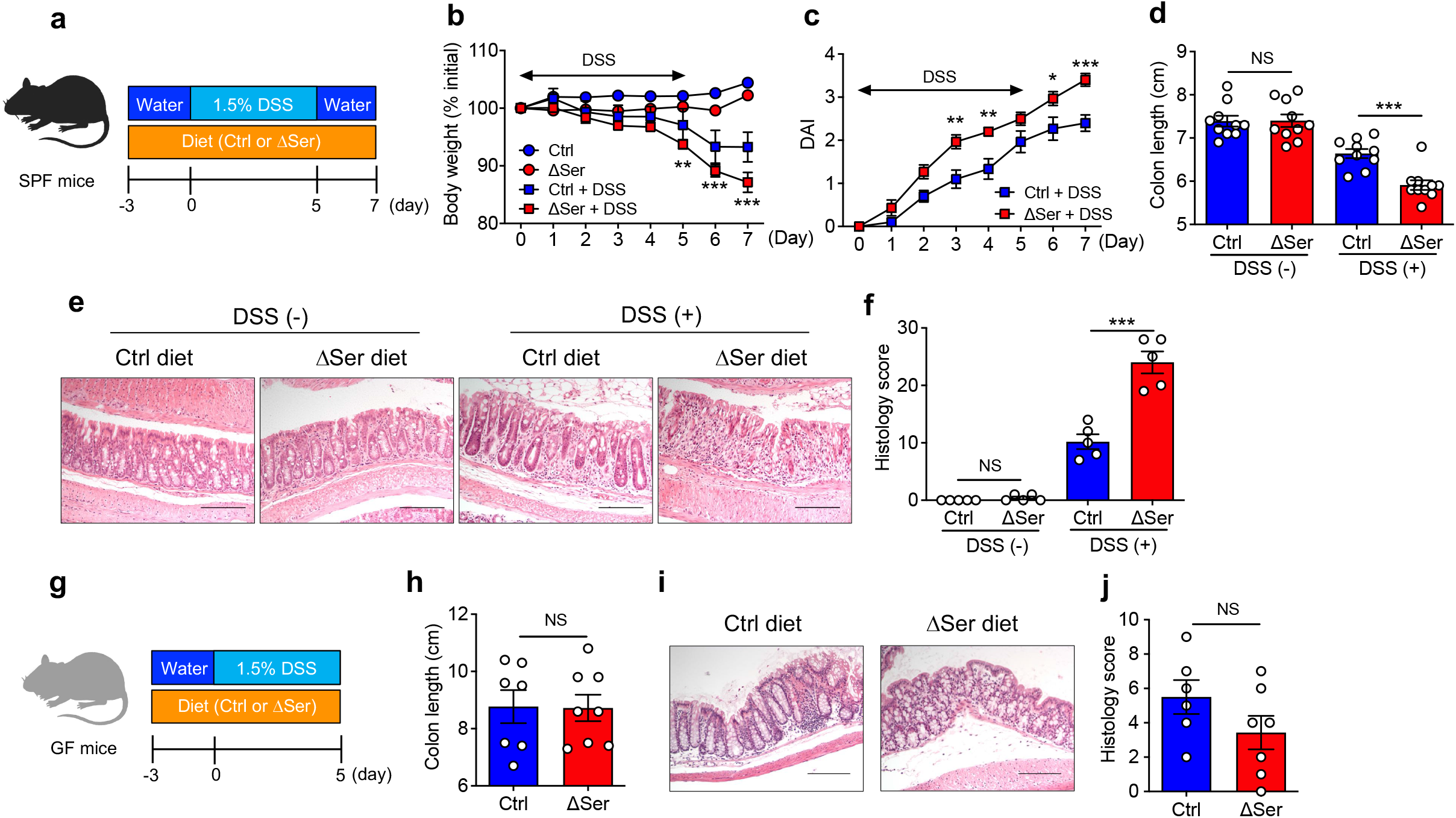
Deprivation of dietary L-serine exacerbates gut inflammation in DSS-induced colitis through the gut microbiota. **a**, SPF C57BL/6 mice were fed the control diet (Ctrl) or the ΔSer diet for 3 days, then given 1.5% DSS for 5 days, followed by conventional water for 2 days. On day 7 post-DSS, all mice were euthanized. **b,c**, Body weight and DAI were monitored during the 5-day DSS treatment. **d–f**, Colon length, representative histological images of colon sections (scale bar, 200 μm), and histology scores were evaluated. **g**, GF Swiss Webster mice were fed a Ctrl diet or a ΔSer diet for 3 days and then treated with 1.5% DSS for 5 days. On day 5 post-DSS, all mice were euthanized. **h–j**, Colon length, representative histological images of colon sections (Scale bar, 200 μm), and histology scores were assessed. Dots indicate individual mice, with mean ± SEM. NS, not significant, \**P* < 0.05, \*\**P* < 0.01, \*\*\**P* < 0.001 by 1-way ANOVA or 2-way ANOVA with Tukey post hoc test or unpaired *t* test. See also **Extended Data Fig. 2**.

### Dietary L-serine starvation leads to the blooms of pathotype *E. coli* in the inflamed gut

We next analyzed the gut microbiota isolated from Ctrl diet– and ΔSer diet–fed mice to identify the bacterial taxa that may be associated with the severe inflammation observed in ΔSer diet–fed mice. We found that the restriction of dietary L-serine affected the microbial composition during inflammation, while in the steady state it had little influence (**Fig. 3a**). Linear discriminant analysis effect size (LEfSe) further identified the bacterial families over- and under-represented after the dietary change. LEfSe analysis showed that Verrucomicrobiaceae and Enterobacteriaceae families were over-represented in ΔSer diet–fed mice, whereas Sutterellaceae and Porphyromonadaceae families were under-represented (**Fig. 3b**). Interestingly, the abundance of *E. coli*, which belongs to the Enterobacteriaceae family, was significantly higher in the colitic mice fed the ΔSer diet rather than the Ctrl diet (**Fig. 3c**). To determine the function of *E. coli* accumulated in ΔSer diet–fed mice, the abundance of genes associated with pathotypes of *E. coli* were evaluated by qPCR. Notably, genes associated with adhesion and invasion to host epithelial cells (vat, fimH, flicC, ompA,, ompC, and ibeA) and metabolic adaptation (pduC, chuA, fyuA) were significantly enriched in ΔSer diet–fed mice than in Ctrl diet–fed mice (**Fig. 3d**). This result suggests that *E. coli* strains accumulated in ΔSer diet–fed mice may be pathotype *E. coli*, such as AIEC. Similarly, the abundance of *Akkermansia muciniphila*, a major bacterial species in the Verrucomicrobiaceae family, was significantly increased in the L-serine deficient condition (**Fig. 3c**). These data indicate that the restriction of dietary L-serine leads to the unexpected blooms of *E. coli* harboring AIEC pathotypes, together with other commensal symbionts, such as *A. muciniphila*, in the inflamed gut.

**Fig 3.**
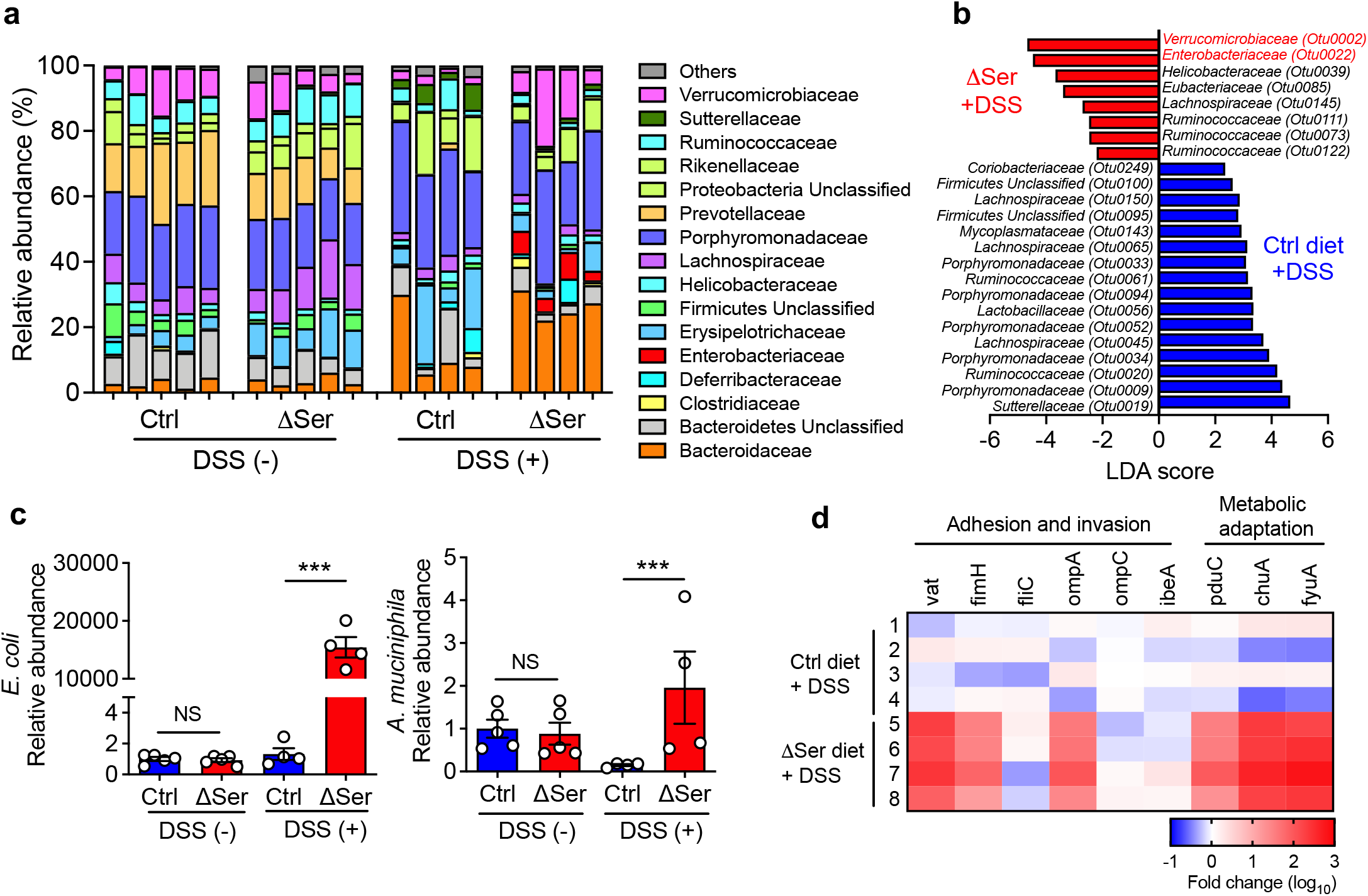
Deprivation of dietary L-serine fosters blooms of pathotype *E. coli* and *A. muciniphila* in the inflamed gut. **a**, Feces were collected from Ctrl diet– and ΔSer diet–fed mice with and without DSS treatment, and DNA was isolated. Gut microbiota was analyzed by 16S rRNA sequencing. **b**, Significantly enriched bacterial taxa in Ctrl diet–fed mice (blue bars) and ΔSer diet–fed mice (red bars) were identified by LEfSe analysis. **c**, The relative abundance of *A. muciniphila* and *E. coli* was each quantified by qPCR. **d**, The heat map shows the abundance of *E. coli* virulence genes in ΔSer diet–fed colitis mice compared to Ctrl diet–fed colitis mice. Dots indicate individual mice, with mean ± SEM. NS, not significant. \*\*\**P* < 0.001 by 1-way ANOVA with Tukey post hoc test.

### *A. muciniphila* enables AIEC to relocate to the epithelial niche by degrading the mucus layer

As AIEC requires L-serine for its fitness in the inflamed gut^4^, we did not expect dietary L-serine restriction to induce an AIEC bloom. The suppression of AIEC growth by dietary L-serine restriction in the setting of intraspecific competition^4^ suggests that other bacterial species in SPF mice may act as metabolic supporters for AIEC to overcome the nutrient limitation. In this regard, we focused on *A. muciniphila* as a metabolic supporter for AIEC. We first assessed the growth kinetics of *A. muciniphila* and *E. coli* during colitis. As colitis progressed, the abundance of *A. muciniphila* gradually decreased in Ctrl diet–fed mice (**Fig. 4a**). In contrast, when dietary L-serine was restricted, *A. muciniphila* was markedly increased on day 1 after DSS treatment and it maintained a higher abundance until day 7 compared (**Fig. 4a**). As *A. muciniphila* does not require L-serine for its growth^26^, it may have a growth advantage over other commensal gut bacteria under the nutrient-restricted condition. Of note, the abundance of *E. coli* (likely enriched with AIEC) was unchanged in the early stage of colitis but dramatically increased at 7 days after DSS treatment with L-serine starvation (**Fig. 4a**). These results suggest that under these conditions, the expansion of *A. muciniphila* may trigger the subsequent proliferation of AIEC.

**Fig 4.**
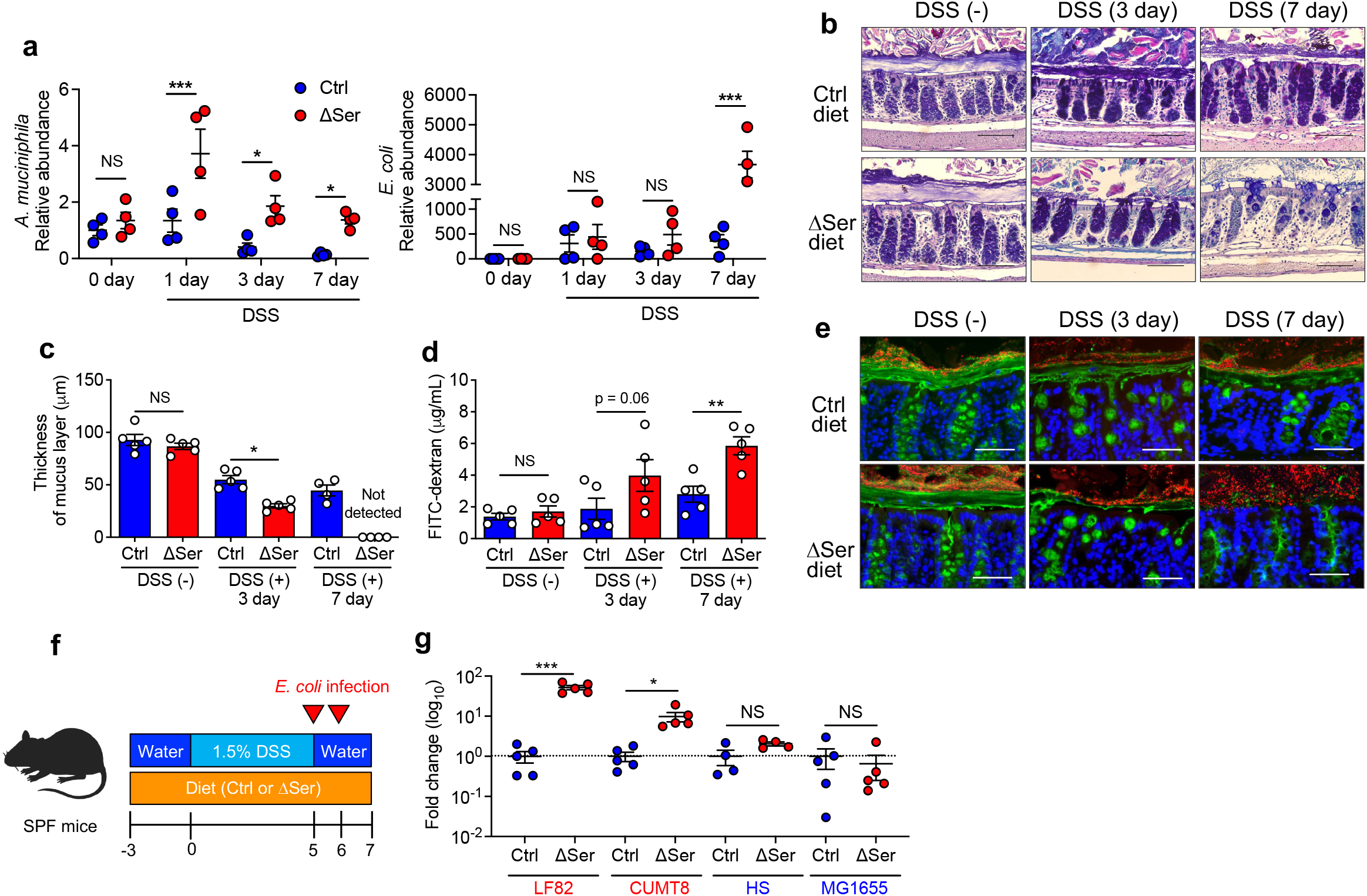
Disruption of colonic mucus barrier under L-serine starvation enhances the encroachment of AIEC to the epithelial niche. **a**, Relative abundance of *A. muciniphila* and *E. coli* during DSS treatment were each assessed by qPCR. **b,c**, Colonic sections were stained with AB/PAS, and the thickness of the inner mucus layer was measured (scale bar, 100 μm). **d**, Intestinal permeability was assessed with FITC–dextran. **e**, Immunostaining (MUC2, green; DAPI, blue) and FISH (EUB338 probe, red,) of Carnoy’s solution–fixed colonic sections (scale bar, 100 μm). **f**, SPF C57BL/6 mice were fed the Ctrl diet or the ΔSer diet for 3 days, then given 1.5% DSS for 5 days, followed by conventional water for 2 days. Mice were infected with each strain of *E. coli* (1 × 10^9^ CFU/mouse) on days 5 and 6. On day 7 post-DSS, all mice were euthanized. **g**, Homogenates of colon tissues were cultured on LB agar plates supplemented with ampicillin or streptomycin. The number of viable bacteria was estimated by counting the CFUs and calculating the fold change (ΔSer diet/Ctrl diet). Dots indicate individual mice, with mean ± SEM. NS, not significant, \**P* < 0.05, \*\**P* < 0.01, \*\*\**P* < 0.001 by 1-way ANOVA with Tukey post hoc test or unpaired *t* test.

*A. muciniphila* is a mucolytic bacterium capable of degrading mucus by several glycoside hydrolases that target the host mucus glycans^26^. Thus, the expansion of *A. muciniphila* may cause mucus barrier dysfunction. Consistent with this notion, we observed that the thickness of the inner mucus layer was significantly reduced in the ΔSer diet–fed mice after DSS treatment, with the expansion of *A. muciniphila* (**Fig. 4b,c**). Notably, in the steady state (i.e., without expanding *A. muciniphila*), the ΔSer diet had no apparent effect on the mucus barrier (**Fig. 4b,c**). As intestinal mucus acts as a physical barrier that keeps luminal antigens, including resident microbiota, distant from the host epithelial cells^27,28^, a defective intestinal mucus barrier may result in the penetration of luminal antigens and increase the risk of colitis^27,29^. Consistent with this notion, dietary L-serine deprivation increased intestinal permeability after DSS treatment (**Fig. 4d**). Also, degradation of the mucus layer by dietary L-serine restriction brought the luminal bacteria closer to the IECs (**Fig. 4e**). Thus, the *A. muciniphila* expansion may promote the encroachment of luminal bacteria, including AIEC, close to the colonic epithelium, and it may contribute to the increased susceptibility to colitis. To validate whether disruption of the mucus barrier under L-serine restriction facilitates the localization of AIEC to the epithelial niche, SPF mice were fed the Ctrl diet or the ΔSer diet to induce the *A. muciniphila* expansion and subsequent mucus barrier disruption in colitic mice (**Fig. 4f**). After disrupting the mucus layer by the feeding of the ΔSer diet, followed by DSS treatment, mice were challenged exogenously with AIEC strains (LF82 and CUMT8) or commensal *E. coli* strains (HS and MG1655) (**Fig 4f**). As shown in **Fig. 4g**, the number of colonic mucosa-associated AIEC strains, but not commensal *E. coli* strains, was significantly higher in the ΔSer diet–fed mice (disrupted mucus layer) than in the Ctrl diet–fed mice (intact mucus layer).

To further examine the direct interaction between *A. muciniphila* and AIEC in the gut, we cocultured *A. muciniphila* and AIEC in the presence of a human-derived colonoid monolayer (HCM). As *A. muciniphila* is an obligate anaerobe^26^, we used a two-chamber system that enables the coculture of anaerobic bacteria in the upper anaerobic chamber and the HCM supplemented with oxygen in the lower aerobic chamber (**Fig. 5a**)^30^. In this culture system, the HCM secreted mucin and formed a thick mucus layer on its surface (**Fig. 5b**). After coculture with *A. muciniphila*, the mucus layer was dramatically reduced (**Fig. 5b**). In contrast, exposure to AIEC strain LF82 did not affect the mucus layer on the HCM (**Fig. 5b**). Notably, the presence of *A. muciniphila* facilitated the encroachment of AIEC LF82 to the HCM (**Fig. 5b**). This consequence was further validated by enumeration of mucosa-associated AIEC. Consistent with the observations of immunofluorescence staining, the association of AIEC LF82 with the HCM was significantly increased in the presence of *A. muciniphila* (**Fig. 5c**). Conversely, the number of nonadherent AIEC LF82 (i.e., floating in media) was even reduced when cocultured with *A. muciniphila* (**Fig. 5c**). These findings suggest that *A. muciniphila*–mediated mucus degradation facilitates the encroachment of AIEC to the epithelial niche.

**Fig 5.**
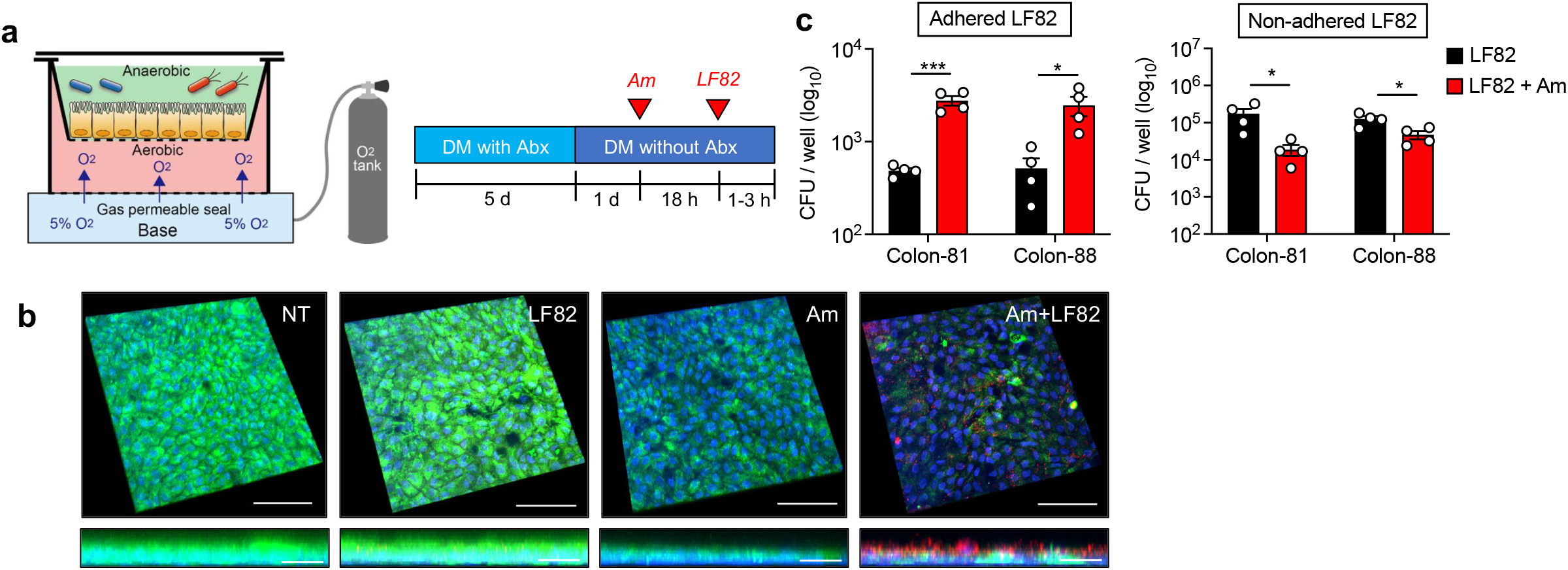
*A. muciniphila*-mediated mucus disruption facilitates adhesion of AIEC to IECs. **a**, Assembly of anaerobic coculture system. The human-derived colonoid monolayer (HCM) from each donor (colon-81 and colon-88) was differentiated for 6 days by differentiation media (DM) with or without antibiotics (Abx). *A. muciniphila* was infected for 18 h, and then AIEC LF82 was infected for 1–3 h. **b**, Immunofluorescence staining of MUC2 (green), *E. coli* (red), and DAPI (blue). Scale bar, 100 μm (XYZ axis) and 20 μm (XZ axis). **c**, Cell-associated AIEC LF82 was cultured on LB agar plates supplemented with ampicillin. The number of viable bacteria was estimated by counting the CFUs. Dots indicate individual mice, with mean ± SEM. N.S., not significant. \**P* < 0.05, \*\*\**P* < 0.001 by unpaired *t* test.

### AIEC and *A. muciniphila* cooperate to promote gut inflammation under dietary L-serine restriction

Thus far, we determined that *A. muciniphila*, which can proliferate independent of L-serine, expands when dietary L-serine is restricted and facilitates the epithelial localization of AIEC strains in the gut by degrading the mucus layer. However, the link between AIEC–*A. muciniphila* interaction and the increased susceptibility to colitis remained unclear. Hence, we next examined the involvement of this bacterial cooperation in the exacerbation of colitis under dietary L-serine restricted conditions. To this end, we generated gnotobiotic mice colonized by AIEC and *A. muciniphila*. To assess the importance of the mucus-degrading capacity of *A. muciniphila*, we used a consortium of nonmucolytic bacterial strains as the base bacterial community. We modified a known synthetic human gut microbiota (SM) model^2^. The original SM consortium is composed of 14 fully sequenced human commensal gut bacteria representing the five dominant phyla, which collectively possess essential core metabolic capabilities^2^. We removed 4 species of mucolytic bacteria (*Bacteroides caccae*, *B. thetaiotaomicron*, *Barnesiella intestinihominis*, and *A. muciniphila*) and commensal *E. coli* from the original consortium. We defined this new base consortium composed of 9 species of nonmucolytic commensal symbionts as the “nonmucolytic synthetic human gut microbiota” (NmSM) (**Fig. 6a**). To evaluate the interaction between AIEC and *A. muciniphila*, we added AIEC LF82 with or without *A. muciniphila* to the base NmSM community (NmSM+LH82 and NmSM+LF82+Am) (**Fig. 6a**). Likewise, for the control groups, the commensal *E. coli* strain HS replaced AIEC LF82 (NmSM+HS and NmSM+ HS+Am) (**Fig. 6a**). All mice were fed with the ΔSer diet, and colitis was induced by DSS treatment (**Fig. 6a**). In the absence of *A. muciniphila*, and under dietary L-serine restriction, the colonization of AIEC LF82 did not exacerbate colitis compared to the colonization of HS(**Fig. 6b–f**). The presence of *A. muciniphila* did not alter the susceptibility to colitis in the commensal *E. coli* HS–colonized mice, although colitis led to a marked expansion of *A. muciniphila* (**Fig. 6b–f**). These results indicated that *A. muciniphila* can gain a growth advantage over other commensal symbionts in the gut when dietary L-serine intake is limited; however, *A. muciniphila* per se is not colitogenic. Notably, unlike commensal *E. coli*, cocolonization of AIEC LF82 and *A. muciniphila* significantly exacerbated colitis (**Fig. 6b–f**). These results suggest that the colonization of AIEC alone is not sufficient to exacerbate colitis, whereas *A. muciniphila* may promote the colitogenic capability of AIEC. Consistent with the severity of colitis, the presence of *A. muciniphila* promoted the expansion of AIEC LF82, but not commensal *E. coli* HS, in both fecal and mucosal compartments (**Fig. 6g–i**). These results demonstrated a causal role of the interaction between AIEC and *A. muciniphila* in the exacerbation of colitis. It is noteworthy that this pathogenic interaction is only observed in the absence of dietary L-serine. In the gnotobiotic mice fed the Ctrl diet (i.e., L-serine–sufficient), the cocolonization of AIEC LF82 and *A. muciniphila* did not increase the susceptibility to colitis (**Extended Data Fig. 3**). The maintenance of the intact mucus layer may explain this phenotype (**Extended Data Fig. 3b,c**).

**Fig 6.**
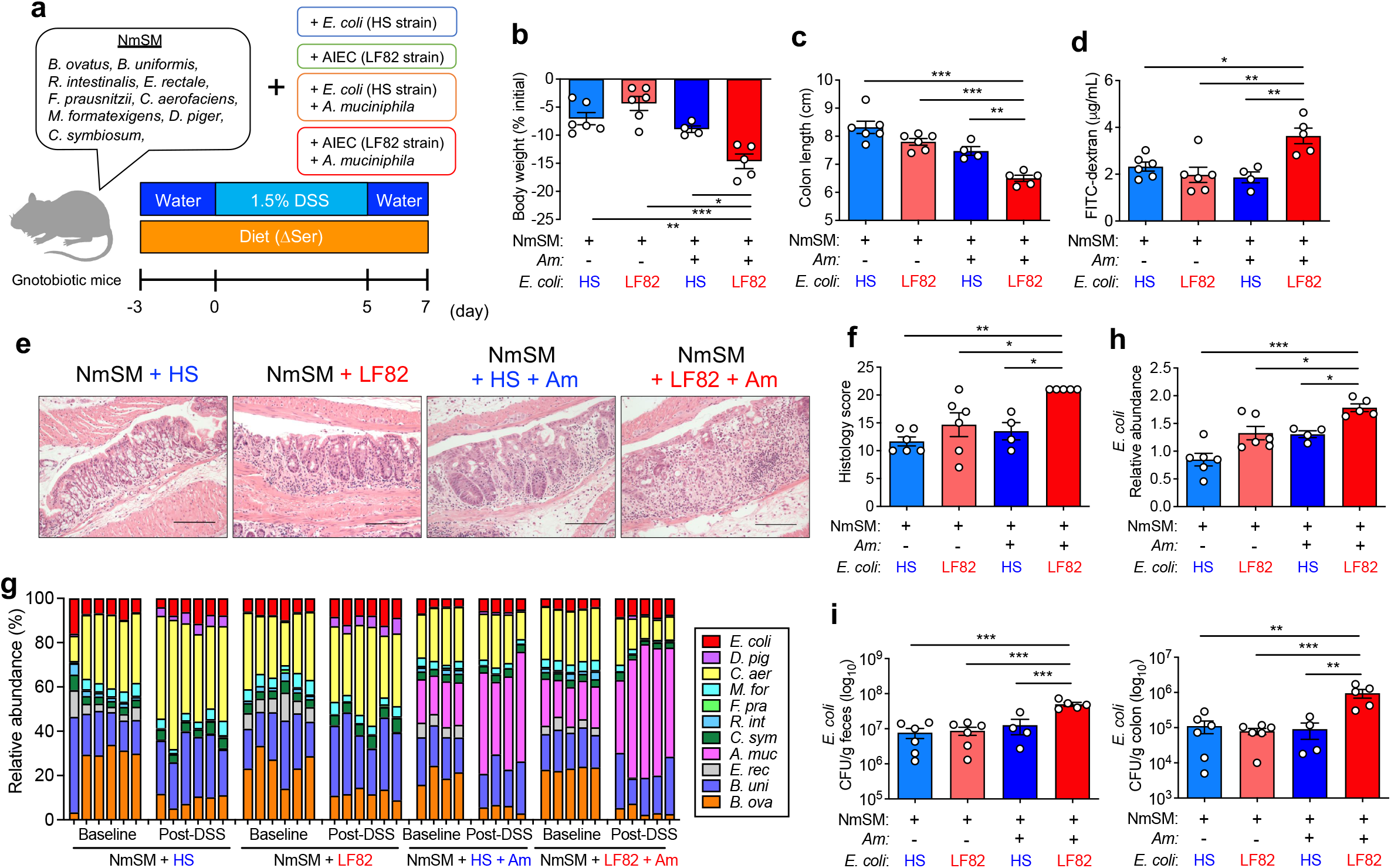
AIEC and *A. muciniphila* cooperatively exacerbate colitis under L-serine restriction. **a**, Experimental protocol and the composition of the nonmucolytic synthetic human gut microbiota (NmSM) for the gnotobiotic mouse experiments. **b,c**, Body weight and colon length were measured 7 days post DSS treatment. **d**, Intestinal permeability was assessed with FITC–dextran. **e,f**, Representative histological images of colonic sections stained with HE (scale bar, 200 μm) and the histology scores. **g,h**, The relative abundance of each bacterial strain at baseline and post-DSS treatment was assessed by qPCR. Fold change of *E. coli* abundance (post-DSS/baseline) was calculated. **i**, Homogenates of feces or colon tissues were cultured on LB agar plates. The number of viable bacteria was estimated by counting the CFUs. Dots indicate individual mice, with mean ± SEM. N.S., not significant. \**P* < 0.05, \*\**P* < 0.01, \*\*\**P* < 0.001 by 1-way ANOVA with Tukey post hoc test. See also **Extended Data Fig. 3**.

### AIEC exploits host-derived L-serine to counteract dietary L-serine deprivation

We demonstrated that mucolytic bacteria promote the relocation of AIEC to the epithelial niche, where it can evade nutrient (i.e., L-serine) restriction, proliferate, and facilitate intestinal inflammation. However, the mechanism by which AIEC proliferates in the epithelial niche under dietary L-serine restriction remained unclear. We hypothesized that AIEC exploits host-derived nutrients in the epithelial niche, as some pathogens can liberate host-derived nutrients for their growth^31^. To determine if AIEC uses host-derived nutrients for its growth, we first compared the growth of LF82 cultured with or without IECs. As shown in **Fig. 7a**, the coculture of AIEC LF82 and IECs significantly enhanced the growth of AIEC LF82 compared with the host cell–free condition. This bacterial growth enhancement by IECs was not observed in the commensal *E. coli* strain HS, nor in the LF82 ΔfimH mutant strain that lacks genes involved in adhesion to IECs (**Fig. 7b**), which suggests that bacterial adhesion to IECs is required for the utilization of host-derived nutrients. To identify the host-derived nutrients used by AIEC in the epithelial niche, we analyzed the transcriptomic changes of AIEC LF82 induced by the association with the HCM (**Extended Data Fig. 4a**). RNA-seq analysis demonstrated that epithelial association significantly altered the transcriptional profiles of AIEC LF82 (**Extended Data Fig. 4b**). The Gene Ontology (GO) enrichment analysis showed that the coculture of AIEC LF82 and HCM upregulated the pathways involved in AIEC growth, including ribosome biosynthesis and protein folding, response to stress, and sugar transport (**Extended Data Fig. 4c**). In contrast, the pathways related to chemotaxis and amino acid biosynthesis were downregulated (**Extended Data Fig. 4c**). In contrast, the presence of AIEC had a minor impact on the transcriptomic profiles of the HCM (**Extended Data Fig. 4d,e**). Notably, AIEC LF82 upregulated genes related to L-serine metabolism and downregulated genes related to L-serine biosynthesis when associated with the HCM (**Fig. 7c**), indicating that AIEC acquires L-serine from the host epithelium and uses it for its growth. In fact, IEC presence did not promote the growth of the Δ*tdc*Δ*sda* (ΔTS) mutant AIEC LF82 strain, which lacks two major L-serine utilization gene operons and is incapable of utilizing L-serine^4^, unlike its effect on LF82 WT (**Fig. 7d**). Consistent with this finding, the association with AIEC LF82 significantly reduced the concentration of free intracellular L-serine in the IECs, whereas the association with LF82 ΔTS had no effect (**Fig. 7e**), indicating that AIEC LF82 consumes L-serine in the infected host cells. Further, we confirmed that although host-derived L-serine is not required for the initial AIEC adhesion and invasion, it is vital for AIEC growth after it associates with the host cells. For example, early phase (1 h) adherence and invasion of AIEC did not differ between the WT and the ΔTS mutant AIEC LF82 strains (**Fig. 7f,g**). However, in the later phase (3 h), proliferation of AIEC after adherence and invasion was significantly impaired in the ΔTS mutant strain compared to the WT strain (**Fig. 7f,g**). Interestingly, the deprivation of L-serine from the media facilitated the adhesion and invasion of AIEC LF82 to IECs (**Fig. 7h**). This evidence suggests that AIEC enhances its growth after adhering to host cells by utilizing host-derived L-serine.

**Fig 7.**
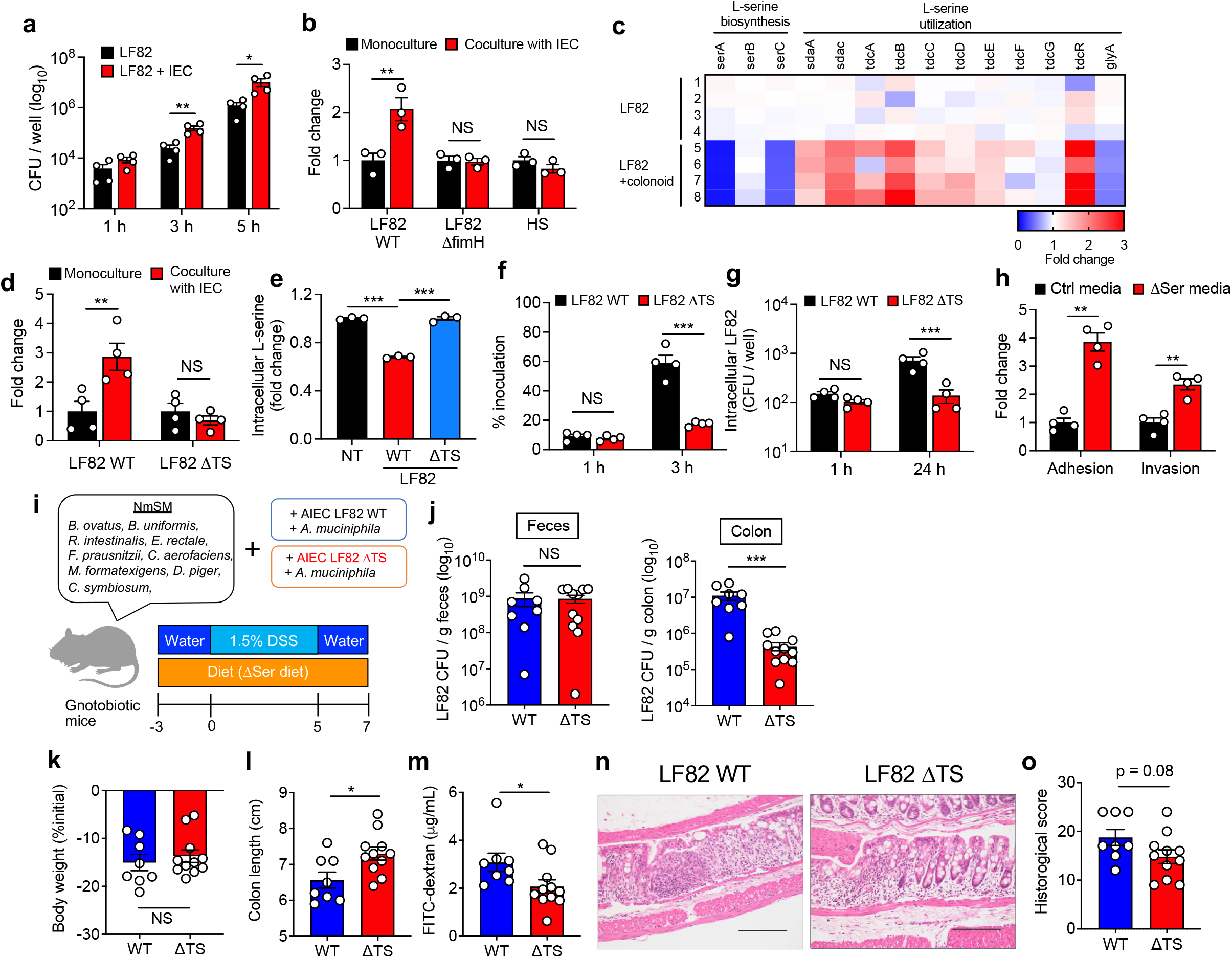
L-serine utilization by AIEC is a partial requirement for the exacerbation of colitis under L-serine deprivation. **a,b**, *E. coli* strains were cultured with and without T84 cells. After 1–5 h infection, the CFUs of total bacteria, including adhered and nonadhered bacteria, were counted. **c**, AIEC LF82 was monocultured or cocultured with the human-derived colonoid monolayer (HCM) for 3 h, and the transcriptomic profiles were evaluated by RNA-seq. The heat map shows fold changes of L-serine metabolism genes (LF82 + HCM/LF82). **d**, LF82 WT or ΔTS mutant strains were infected in T84 cells for 5 h, and then the CFUs of total bacteria were counted. **e**, Fold changes of intracellular L-serine after infection of T84 cells with AIEC strain LF82. **f**, T84 cells were infected with LF82 WT or ΔTS mutant strains. After 3 h, adhesion bacteria were counted. **g**, T84 cells were infected with LF82 WT or ΔTS mutant strains. After 1 h, the cells were cultured with gentamicin (100 μg/mL) for 24 h. Intracellular bacteria were plated on LB agar plates and counted. **h**, LF82 and T84 cells were cocultured in the Ctrl media or ΔSer media. After a 3 h infection, adhesion and invasion bacteria were plated on LB agar plates and counted. **I**, Experimental design. GF mice were colonized by nonmucolytic synthetic human gut microbiota (NmSM) and *A. muciniphila* with LF82 WT or ΔTS mutant strains. **j**, On day 7 post-DSS, all mice were euthanized, and the LF82 burden in the colon and feces was assessed. **k,l**, Body weight and colon length. **m**, Intestinal permeability was evaluated by FITC–dextran assay. **n,o**, Representative histological images (scale bar, 200 μm) and histology scores were evaluated. Dots indicate individual mice, with mean ± SEM. N.S., not significant. \**P* < 0.05, \*\**P* < 0.01, \*\*\**P* < 0.001 by 1-way ANOVA with Tukey post hoc test or unpaired *t* test. See also **Extended Data Fig. 4**.

Lastly, we assessed the extent to which the utilization of host-derived L-serine by AIEC is linked to the susceptibility to colitis. To this end, NmSM-colonized gnotobiotic mice were colonized with either AIEC LF82 WT or ΔTS mutant AIEC LF82 together with *A. muciniphila* (**Fig. 7i**). Both groups of mice were fed the ΔSer diet, followed by a DSS challenge. The colonization of LF82 WT or the ΔTS mutant was comparable in feces, whereas the number of AIEC associated with the colonic mucosa was significantly lower in the ΔTS mutant strain than in the LF82 WT strain (**Fig. 7j**), suggesting that AIEC LF82 may exploit host-derived L-serine in the epithelial niche. Furthermore, consistent with the impaired proliferation of AIEC in the epithelial niche, the mice colonized with the LF82 ΔTS mutant strain displayed an attenuated degree of colitis compared to the mice colonized with the LF82 WT strain (**Fig. 7k–o**).

## DISCUSSION

In this study, we show that the biosynthesis and utilization of L-serine are disturbed in the gut microbiota of patients with IBD, which is consistent with our recent research showing that the IBD-associated pathobiont AIEC upregulates L-serine catabolism in the inflamed gut^4^. We had expected that the deprivation of dietary L-serine would attenuate inflammation by suppressing the expansion of pathobionts, such as AIEC. However, we observed that dietary L-serine restriction leads to the expansion of AIEC and subsequent exacerbation of colitis. This unexpected and adverse effect of dietary L-serine deprivation is context dependent. Dietary L-serine restriction promotes the abnormal expansion of AIEC only when it coexists with mucolytic bacteria, such as *A. muciniphila*. The fact that L-serine is a crucial nutrient for the growth of various gut bacteria, including AIEC, but not *A. muciniphila*, gives *A. muciniphila* a growth advantage over other commensal microbes under L-serine restriction. Notably, the blooms of *A. muciniphila* per se are not detrimental. However, amassed *A. muciniphila* facilitates the encroachment of AIEC to the epithelial niche by degrading the mucus barrier. As a result, AIEC can reside in the epithelial niche and counteract dietary L-serine restriction by extracting host-derived nutrients. Notably, AIEC can acquire L-serine pooled in the host colonic epithelium. This novel insight advances the understanding of the complex interplay among pathobionts, symbionts, and host cells in the context of gastrointestinal diseases, such as IBD.

Dietary amino acids are vital nutrients for maintaining intestinal homeostasis and the gut microbiota^32^. L-serine is thought to be a conditionally essential amino acid as it plays a critical role in the cellular metabolisms of both mammalian and bacterial cells only under certain conditions^4,19-21^. Several biosynthetic and signaling pathways require L-serine, including the synthesis of other amino acids, the production of phospholipids, and the provision of one-carbon units to the folate cycle, which are used for the de novo synthesis of nucleotides^33^. L-serine also contributes to the production of glutathione, which is essential for reducing toxic oxidants and metabolic byproducts. Thus, L-serine supports several metabolic processes that are crucial for growth and adaptation to the microenvironment. It has been reported that some *E. coli* strains rapidly consume L-serine, as compared to other amino acids^34^. Notably, *E. coli* promotes the consumption of L-serine under heat stress^35^. Consistently, our work has demonstrated that bacteria belonging to the Enterobacteriaceae family, including *E. coli*, use L-serine to adapt to the inflamed gut^4^. Thus, some bacteria, most likely pathogens, utilize L-serine metabolism to counteract environmental stress. The multi’omics database shows that L-serine metabolism is disturbed in parallel with a lower fecal L-serine concentration in IBD. Although we need to clarify the mechanism by which gut bacteria consume L-serine in IBD, certain bacteria may use L-serine to adapt to the inflammatory microenvironment. How the concentration of luminal L-serine is regulated remains unclear. Diet is the primary source of luminal L-serine, as dietary deprivation significantly reduces its concentration in the gut lumen^4^. Regarding the consumption of L-serine, both host cells and bacteria are L-serine utilizers in the gut^19,36,37^. Generally, L-serine is contained in protein-rich foods, such as meat, fish, eggs, and soybeans. A recent systematic review has shown that fiber and calcium intakes are insufficient in IBD, whereas protein intake meets or exceeds the recommended amount^38^. Therefore, the reduced concentration of L-serine in the gut lumen of individuals with IBD may be caused by the excessive consumption of L-serine by the gut microbiota or the host cells rather than by the insufficient intake of protein. As mentioned, L-serine plays a pivotal role in the fitness of certain bacteria, including pathobionts such as AIEC, particularly when exposed to environmental stress. Consequently, pathobionts may evolve backup systems to evade the shortage of such a vital nutrient. This seems to be a strategy that pathobionts use to maintain a fitness advantage over commensal competitors in disease conditions. In the present study, we found that AIEC can switch the source of L-serine from the diet to the host cells when dietary L-serine is limited. To acquire host-derived L-serine, AIEC relocates its colonizing niche from the gut lumen to the mucosa. Given the evidence that the availability of L-serine in the gut lumen is decreased in patients with IBD, the enrichment of mucosa-associated AIEC in IBD may be triggered by the perturbed amino acid homeostasis in the gut lumen.

As L-serine metabolism plays a vital role in the survival of some gut microbes, particularly pathobionts, this metabolic pathway can be a therapeutic target for pathobiont-driven inflammatory diseases, such as IBD. Our previous work showed that the dietary deprivation of L-serine suppresses AIEC expansion^4^. In the current study, dietary L-serine deprivation resulted in the unexpected expansion of AIEC. Thus, the therapeutic effects of the dietary intervention are context dependent. In other words, the same dietary therapy could be either beneficial or detrimental. The factor that dictates the efficacy of dietary interventions may be the basal microbiota composition of individuals. For example, dietary L-serine restriction may not significantly impact patients who are not colonized by AIEC or colonized by AIEC without coexistent mucolytic bacteria. In individuals colonized by AIEC and metabolic competitors for AIEC (e.g., certain commensal *E. coli* strains) but lacking mucolytic bacteria, dietary L-serine restriction can facilitate the competitive elimination of AIEC by the commensal *E. coli* strains. Indeed, gnotobiotic mice colonized by the microbiota from a patient with CD displayed a reduction of Enterobacteriaceae and the attenuation of colitis when L-serine was removed from the diet ^4^. This result implies that this individual had a sufficient number of metabolic competitors for AIEC and the restriction of dietary L-serine prompted the competitive elimination of AIEC. On the other hand, as shown in the current study, in the presence of mucolytic bacteria, AIEC can overcome nutritional restriction and exacerbate colitis. Notably, *A. muciniphila* has a protective role in some individuals with metabolic diseases, such as obesity and diabetes mellitus^39,40^, and therefore, it has been proposed as a probiotic bacterium^41^. Consistently, in the present study, *A. muciniphila* colonization per se did not exacerbate colitis, even under dietary L-serine restriction (with a noticeable *A. muciniphila* expansion). However, *A. muciniphila* can serve as a metabolic supporter for AIEC, and hence, indirectly contribute to the pathogenesis of colitis. Thus, the balance of pathobionts and their metabolic competitors and supporters may determine the outcomes of dietary interventions.

Nutritional competition is one of the main strategies used by commensal symbionts to prevent the colonization and proliferation of commensal pathobionts and exogenous pathogens^42,43^. To overcome this symbiont-mediated colonization resistance, pathogens have evolved various strategies^44^. For example, some pathogens use unique nutrients, such as ethanolamine, that commensal symbionts cannot use^45^. Likewise, relocation of the living niche provides an escape from the nutritional competition. For example, *Citrobacter rodentium*, a mouse pathogen used to model human infections with enteropathogenic *E. coli* (EPEC) and enterohemorrhagic *E. coli* (EHEC), resides on the intestinal epithelial surface by expressing the locus of enterocyte effacement (LEE) virulence factor. In this new niche, *C. rodentium* can acquire specific nutrients not accessible to commensal symbionts residing in the luminal niche^46^. However, unlike *C. rodentium* or other enteropathogens, commensal pathobionts, such as AIEC, lack the LEE virulence factors, and therefore the niche relocation is rarely executed in the steady-state (i.e., intact gut microbiota and mucus barrier). These pathobionts are, therefore, not classified as obligate pathogens, but rather as opportunistic pathogens. In other words, pathobionts have a poor ability to proliferate and establish infection in the healthy hosts. Instead, pathobionts bloom only in conditions that compromise the host’s defenses, including the colonization resistance by the healthy gut microbiota. In this context, we found that mucolytic bacteria, such as *A. muciniphila*, can serve as metabolic supporters for AIEC in the gut. Although *A. muciniphila* may not directly promote the growth of AIEC, *A. muciniphila* licenses AIEC to acquire alternative nutrients by facilitating its niche relocation.

Some pathogens, like AIEC, which can reside in the epithelial niche, can obtain nutrients from infected host cells for replication. For example, EPEC exploits nutrients from infected host cells by using injectisome components, which enable the pathogen to thrive in competitive niches^47^. As AIEC strains lack injectisome components, it is plausible that AIEC exploits a distinct mechanism to extract host-derived nutrients when associated with the epithelial niche. In this regard, AIEC may invade the colonic epithelium to acquire nutrients from the host cells rather than extracting nutrients from the cell surface, as cellular invasion is a unique pathogenic feature of AIEC strains compared to other pathogenic *E. coli* strains^48,49^. Further research is required to identify the mechanisms by which AIEC exploits host-derived nutrients. Our present study shows that AIEC uses L-serine pooled in the host epithelium. This notion was supported by the upregulation of L-serine utilization genes in AIEC when associated with the IECs, along with the reduced free L-serine concentration in the IECs. Moreover, the growth promotion of AIEC due to the AIEC–IEC interaction does not occur if the AIEC strain is incapable of utilizing L-serine (i.e., ΔTS mutant AIEC LF82). Along with the aforementioned critical roles of L-serine that enable AIEC to adapt to the inflammatory environment, our results confirm that L-serine is a major nutrient extracted from the host cells by AIEC in this setting. However, AIEC may extract and use other host-derived nutrients in addition to L-serine. In this context, several pathogens exploit host glycan metabolites as nutrient sources. For example, *Salmonella enterica* serovar Typhimurium and *Clostridioides difficile* use sialic acid liberated from host mucus glycans by *Bacteroides thetaiotaomicron*^50^. EHEC uses fucose, also liberated from host glycans by *B. thetaiotaomicron*, for the regulation of virulence factor expression and colonization in the gut^51^. AIEC strains can also use fucose as a nutrient source through propanediol dehydratase^52,53^. Consistent with this notion, AIEC upregulated genes related to the metabolism of other possible host-derived nutrients (e.g., mannose, *N*-acetylglucosamine) when associated with IECs. These other metabolites may compensate for the growth of AIEC, at least to some extent, in the absence of L-serine. Nevertheless, the studies by us and others as described suggest that L-serine plays a major role among diet- and host-derived nutrients accessible to AIEC in regulating its fitness.

Considered in their entirety, our results demonstrate that the pathogenic capacity of commensal pathobionts, such as AIEC, is context dependent. In the steady-state gut, pathobionts may behave as nonpathogenic commensals. Pathobionts may be detrimental only when metabolic supporters are present. Notably, the metabolic supporters, such as mucolytic bacteria, per se are not detrimental. Also, the balance of luminal nutrients, regulated by diet, is essential to elicit the interaction between pathobionts and metabolic supporters. Therefore, the complex pathobiont–symbiont interactions dictate the success of dietary interventions. Hence, a personalized dietary intervention adapted to the composition of an individual’s gut microbiota is required to treat IBD effectively.

## Method

### Human data

Metagenomics, metatranscriptomics, and metabolomics data were downloaded from the public resource, the second phase of the Integrative Human Microbiome Project (HMP2 or iHMP) – the Inflammatory Bowel Disease Multi’omics Database (https://ibdmdb.org/). The description and collection of samples, and the data preprocessing are explained in a previous study^22^. The samples included in the current analysis are described in **Fig. 1a**.

### Animal

SPF C57BL/6 mice were housed by the Unit for Laboratory Animal Medicine at the University of Michigan. GF Swiss Webster mice were obtained from the Germ-Free Mouse Facility at the University of Michigan. GF mice were housed in flexible film isolators, and their germ-free status was confirmed weekly by aerobic and anaerobic culture. Female and male mice, age 6 – 12 wk, were used in all experiments. Mice were fed either a control amino acid–based diet (Ctrl., TD.130595) or an L-serine deficient diet (ΔSer, TD.140546), which had been used previously^4,18^. The custom diets were manufactured by Envigo (Madison, WI) and sterilized by gamma irradiation. All animal studies were performed in accordance with protocols reviewed and approved by the Institutional Animal Care and Use Committee at the University of Michigan.

### Histology

The colon tissues were quickly removed and immediately preserved overnight in 4% paraformaldehyde for regular histology assessment or in Carnoy’s fixative (60% dry methanol, 30% chloroform, 10% glacial acetic acid) for mucus barrier evaluation. The preserved colon tissues were then incubated in 70% ethanol or dry methanol, respectively, and processed into paraffin-embedded tissue sections (4–5 μm) and stained with HE for histological assessment. Histological inflammation was scored at the In-Vivo Animal Core in the Unit for Laboratory Animal Medicine at the University of Michigan. A veterinary pathologist performed a blind evaluation of the histological scores. Inflammation and epithelial loss were assessed for severity based on the most severe lesion in each section (i.e., 0, none; 1, mild; 2, moderate; 3, severe; 4, marked). Lesion extent was assessed as the percent of the section affected (0, 0%; 1, 1%–25%; 2, 26%–50%; 3, 51%–75%; 4, 76%–100%). The extent and severity scores for inflammation and epithelial cell loss were multiplied to give a total score for each parameter (range 0–16). The total scores for each parameter were summed to give a total colitis score (range 0–32).

### DNA extraction, qPCR, and 16S rRNA sequencing

Fecal DNA was extracted using the DNeasy Blood and Tissue Kit (Qiagen, Germantown, MD), according to a procedure used in a previous study^36^. In the gnotobiotic experiments, fecal DNA was isolated by phenol:chloroform:isoamyl alcohol, as previously described^54^. Briefly, 500 μL buffer A (200 mM NaCl, 200 mM Tris, 20 mM EDTA), 210 μL 20% SDS (filter sterilized), and 500 μL phenol:chloroform:isoamyl alcohol (125:24:1, pH 8.0, Thermo Fischer Scientific, Waltham, MA) were added to the cecal content. The mixture was subjected to bead beating for 3 min at 4°C and centrifuged at 4°C (17,000*g* for 3 min). The aqueous phase was recovered, and 500 μL phenol:chloroform:isoamyl alcohol (pH 8.0) was added. After mixing with a vortex mixer, the mixture was centrifuged again at 4°C (17,000*g* for 3 min). The aqueous phase was recovered and 500 μL of chloroform was added and mixed by inversion. After centrifuging (17,000*g*) for 3 min, the aqueous phase was transferred to new tubes, and 60 μL 3 M sodium acetate (pH 5.2) and 600 μL isopropanol were added. After incubation for 60 min at −20°C, the mixture was centrifuged at 4°C (13,000 rpm for 20 min). The pellet was washed with 70% ethanol, and resuspended in nuclease-free water. DNA was further cleaned using the DNeasy Blood and Tissue Kit (Qiagen). qPCR was performed using a Radiant SYBR Green Lo-ROX qPCR Kit (Alkali Scientific, Fort Lauderdale, FL). To eliminate the inhibitory effect of dextran sodium sulfate (DSS) on qPCR, spermine was added to the PCR ^55^. The relative expression of the target bacteria was calculated using universal 16S primers as a reference. In the gnotobiotic experiments, the relative abundance of bacteria was calculated using the standard curves obtained from the monoculture of each strain^54^. The primer sets used for amplification are listed in **Supplementary Table 1**. For the 16S rRNA sequencing, PCR and library preparation were performed at the Microbiome Core at the University of Michigan. The V4 region of the 16S rRNA-encoding gene was amplified from extracted DNA using the barcoded dual-index primers, as reported previously^56^. Samples were amplified, normalized, and sequenced on the MiSeq system. Raw sequences were analyzed using mothur (v1.33.3). Operational taxonomic units (OTUs) (>97% identity) were curated and converted to relative abundance using mothur. We performed LEfSe to identify significant differentially abundant OTUs.

### Measurement of the thickness of the colonic mucus layer

To measure the thickness of the colonic inner mucus layer, the colonic sections were stained with alcian blue/periodic acid–Schiff (AB/PAS) according to the following protocol: 1) deparaffinize and hydrate in distilled water, 2) 3% acetic acid for 3 min, 3) AB solution for 15 min, 4) wash in running tap water for 2 min, 4) periodic acid solution for 5 min, 5) rinse with distilled water, 6) Schiff reagent for 15 min, 7) wash in running tap water for 5 min, 8) Gill hematoxylin solution for 90 sec, 9) wash in running tap water for 5 min, 10) dehydrate and clear in xylene, 11) cover with a coverslip. We used the images captured of all the available fecal masses of each mouse for quantification, although this number was variable, and generally, the colitic mice had fewer colonic fecal masses. The thickness of the mucus layer in the colonic sections was measured using ImageJ.

### FISH and immunofluorescence staining

FISH and MUC2 immunofluorescence staining were performed according to a previous study, with slight modifications^57^. Briefly, paraffin-embedded colon sections were deparaffinized and hydrated. Sections were then incubated with 2 μg Alexa555-conjugated EUB338 (5′-GCTGCCTCCCGTAGGAGT-3′) in 200 μL hybridization buffer (20 mM Tris-HCl, 0.9 M NaCl, 0.1% (w/v) SDS) at 50°C. After overnight incubation, sections were rinsed in wash buffer (20 mM Tris-HCl, 0.9 M NaCl) at 50°C for 15 min. Sections were blocked with 1% BSA in PBS at room temperature for 1 h and then incubated with anti-MUC2 antibody (H-300; Santa Cruz Biotechnology, Dallas, TX) at 4°C for 6 h. After washing with PBS, sections were incubated with Alexa 488–conjugated rabbit polyclonal antibody (Invitrogen, Thermo Fisher Scientific, Waltham, MA) and DAPI at room temperature for 1 h. To reduce autofluorescence, the sections were treated with an autofluorescence quenching kit (Vector Laboratories, Burlingame, CA), according to the manufacturer’s instruction. The slides were stored overnight, in the dark, at room temperature, and then visualized using the Nikon Eclipse TE2000-S inverted microscope (Nikon USA, Melville, NY). For the in vitro experiment, immunofluorescence staining of MUC2 and *E. coli* was performed as previously described, with slight modifications^58^. Briefly, the human-derived colonoid monolayer (HCM) was fixed with Carnoy’s solution (90% dry methanol and 10% glacial acetic acid), washed 2 times with PBS, permeabilized with 0.1% Triton-X for 10 min, and blocked with 1% BSA/PBS for 30 min. Cells were then incubated with MUC2 antibody (H-300; Santa Cruz Biotechnology) and anti–*E. coli* LPS antibody (2D7/1; Abcam, Waltham, MA) in 1% BSA/PBS overnight at 4°C. The cells were then washed with PBS twice for 5 min followed by incubation with Alexa Fluor 488–conjugated goat anti-rabbit antibody (Invitrogen), Alexa Fluor 555–conjugated goat anti-mouse antibody (Invitrogen), and DAPI for 1 h at room temperature. Stained cells were analyzed using a Nikon A1 confocal microscope.

### Intestinal permeability assay

Intestinal permeability assay was performed using FITC–dextran, as described previously^59^. Briefly, mice were deprived of food for 4 h and then gavaged with 0.6 mg/g body weight 4 kDa FITC–dextran (FD4, Sigma-Aldrich St. Louis, MO). Blood was collected after 4 h, and fluorescence intensity was measured (excitation, 485 nm; emission, 520 nm). FITC–dextran concentrations were determined using a standard curve generated by the serial dilution of FITC–dextran.

### Gnotobiotic experiments

GF Swiss Webster mice were colonized by a consortium of human synthetic microbiota, as reported previously^2^, with a few modifications. The composition of the bacteria consortium is shown in **Fig. 6a and Extended Data Fig. 3a**. Bacteria were anaerobically (i.e., 85% N_2_, 10% H_2_, 5% CO_2_) cultured in their respective media at 37°C with final absorbance (600 nm) readings ranging from about 1.0. *Bacteroides ovatus* (DSMZ 1896), *B. uniformis* (ATCC 8492), *Clostridium symbiosum* (DSMZ 934), *Collinsella aerofaciens* (DSMZ 3979), and *E. coli* (ATCC HS and CD patient–derived LF82) were cultured in TYG medium^60^. Modified chopped meat medium^61^ was used to culture *A. muciniphila* (DSMZ 22959) and *Eubacterium rectale* (DSMZ 17629). *Roseburia intestinalis* (DSMZ 14610), *Faecalibacterium prausnitzii* (DSMZ 17677), and *Marvinbryantia formatexigens* (DSMZ 14469) were cultured in YCFA medium^62^. *Desulfovibrio piger* (ATCC 29098) was cultured in ATCC 1249 medium. Bacterial cultures were mixed in equal volumes, and the mice were orally gavaged with 0.2 mL of this mixture. After a 14-day reconstitution, the mice were fed a sterilized custom diet and treated with DSS.

### Cell culture

T84 cells derived from a human colorectal carcinoma cell line were purchased from ATCC (Gaithersburg, MD) and cultured in Ham’s F-12 nutrient mixture + DMEM (1:1) supplemented with 10% FBS and an antibiotic solution (penicillin-streptomycin). The human stem cell–derived colonoid line was cultured as described in the protocol from the Translational Tissue Modeling Laboratory at the University of Michigan (https://www.umichttml.org). Histologically normal colon tissue from subjects (donors 81, 84, and 88) in our previous studies was used for the colonoid culture^63,64^. The collection and use of human colonic tissue for the colonoids were approved by the Institutional Review Boards (IRBMED) at the University of Michigan. The experiment was conducted according to the principles stated in the Declaration of Helsinki.

### *E. coli* infection in vivo and in vitro

For *E. coli* infection in vivo, mice were infected with each *E. coli* strain (1 × 10^9^ colony forming units (CFU)/mouse). To assess *E. coli* colonization, homogenates of feces and colon tissues were cultured on LB agar plates with ampicillin or streptomycin. The number of viable bacteria was estimated by plate counting the number of CFUs. In the in vitro experiments, T84 cells or the HCM were infected with *E. coli* at an MOI = 1–10 (2 × 10^5^–2 × 10^6^ CFU/well) as described in the respective **Fig. 5, 7, and Extended Data Fig. 4**. After infection, the cells were centrifuged at 1,000*g* for 10 min at 24°C and maintained at 37°C. In the bacterial adhesion assay, cells were washed three times with PBS and then lysed with 0.1% Triton X-100 (Sigma-Aldrich) in deionized water. In the bacterial invasion assay, cells were infected with *E. coli* for 1–3 h and then cultured with gentamycin (100 μg/mL) to kill extracellular *E. coli*. After incubation, the cells were washed 3 times with PBS and then lysed with 0.1% Triton X-100 in deionized water. Lysed cells were diluted and plated on LB agar plates to determine the number of CFUs corresponding to the total number of cell-associated bacteria.

### Coculture of anaerobic bacteria with primary human colon monolayers

The human-derived colonoids were provided by the Translational Tissue Modeling Laboratory at the University of Michigan. To generate the monolayers, the three-dimensional human colonoids were dissociated into a single-cell suspension and plated on collagen IV (Sigma-Aldrich)–coated transwell inserts (0.4 μm pore size, 0.33 cm^2^, polyester [PET], Costar, Corning, Tewksbury, MA) in a 24-well plate. After 24 h, the growth medium was replaced with differentiation medium^65^. Differentiation was completed in a 5% O_2_, 5% CO_2_, balanced N_2_ environment. The medium was refreshed every 48 h for 6 days. Transepithelial electrical resistance (TEER) was determined to be present in an intact monolayer (>400 Ω/cm^2^), which is suitable for an anaerobic coculture system. TEER was recorded using an electrical volt/ohm meter (World Precision Instruments, Sarasota, FL). The transwell plates were set in the apical chambers (Coy Laboratory Products, Grass Lake, MI) of an anaerobic chamber (5% CO_2_, balance N_2_ environment), and cultured in an anaerobic coculture system as shown in **Fig. 6a**^30^. Briefly, setting up the apical chamber, a 24-well gas-permeable plate was placed on the base and sealed in place using double-sided adhesive tape. The entire apparatus was placed in an anaerobic chamber (90% N_2_, 5% H_2_, 5% CO_2_) to allow the growth of anaerobic bacteria in the apical wells. 5.0% O_2_ was pumped from an external tank through the base of the plate to supply oxygen to the basolateral side of the monolayer. The basal compartment was maintained in a 5% O_2_, 5% CO_2_, balance N_2_ environment, whereas the apical chamber was maintained in a 5% CO_2_, balance N_2_ environment. The colonoid monolayer was cultured with differentiation medium in the basolateral wells and minimal coculture medium in the apical wells. To evaluate the impact of *A. muciniphila* on the integrity of the mucus layer and the adhesion of AIEC, the apical well was infected with *A. muciniphila* (4 × 10^4^ CFU/well) for 18 h, and then with LF82 (4 × 10^3^ CFU/well) for 3 h. LF82 adhesion was assessed as described.

### Bacterial RNA-sequencing

Bacterial RNA was isolated with a bacterial RNA isolation kit (Omega Bio-tek, Norcross, GA), according to the manufacturer’s protocol. Isolated RNA was treated with DNase (Thermo Fisher Scientific) and then cleaned with an RNeasy Mini Kit (QIAGEN). Library preparation and sequencing of the RNA-seq libraries were performed in the Advanced Genomics Core at the University of Michigan. Briefly, RNA was assessed for quality using the Agilent TapeStation system (Agilent, Santa Clara, CA). Samples were prepared using the New England BioLabs (Ipswitch, MA) NEBNext Ultra II Directional RNA Library Prep Kit for Illumina, the NEBNext rRNA Depletion Kit (Bacteria), the NEBNext rRNA Depletion Kit (Human/Mouse/Rat), and the NEBNext Multiplex Oligos for Illumina (Unique Dual Index for Primer Pairs), where 145 ng total RNA was ribosomal depleted using the human/mouse/rat and bacteria rRNA depletion modules. The rRNA-depleted RNA was then fragmented for 7–10 min determined by the RIN (RNA Integrity Number) of input RNA as per protocol and copied into first-strand cDNA using reverse transcriptase and dUTP mix. The samples underwent end repair and dA-tailing, followed by ligation of the NEBNext adapters. The products were purified and enriched by PCR to create the final cDNA libraries, which were checked for quality and quantity by the Agilent TapeStation system and Qubit (Thermo Fisher Scientific). The samples were pooled and sequenced on the Illumina NovaSeq 6000 S4 paired-end 150 bp, according to the manufacturer’s recommended protocols (Illumina, San Diego, CA. bcl2fastq2 Conversion Software (Illumina) was used to generate de-multiplexed Fastq files. Paired-end reads were mapped on LF82 genomic sequence CU651637 by bowtie2 and reads of individual LFL82 genes were counted by HTSeq as described^4^. Human transcripts were mapped on human cDNA sequences GRCg38 and counted by Salmon (https://pubmed.ncbi.nlm.nih.gov/28263959/). For functional GO enrichment analysis, over-represented and under-represented bacterial genes were identified by LEfSe and then analyzed using the Gene Ontology Resource (http://geneontology.org)

### Statistical analysis

Statistical analyses were performed using GraphPad Prism 9.3.0 (GraphPad Software San Diego, CA). The numbers of animals used for individual experiments, details of the statistical tests used, and pooled values for several biological replicates are indicated in the respective figure legends. Differences between the two groups were evaluated using the two-tailed Student *t* test or the Mann–Whitney U test. 1-way ANOVA or the Kruskal–Wallis test followed by the Tukey correction or the Dunn test was performed for the comparison of more than 3 groups. Differences of p⎕<⎕0.05 were considered significant. Statistically significant differences are shown with asterisks as follows: *p < 0.05, **p < 0.01, ***p < 0.001, whereas N.S. indicates comparisons that are not significant.

### Data availability

The accession number for the 16S rRNA MiSeq data and bacterial RNA sequence data reported in this paper is PRJNA763203.

## Acknowledgements

The authors wish to thank the University of Michigan Center for Gastrointestinal Research (NIH 5P30DK034933), and the Host Microbiome Initiative, the Germ-Free Mouse Facility, the In-Vivo Animal Core, the Advanced Genomics Core, the School of Dentistry Histology Core, the Rogel Cancer Center Tissue & Molecular Pathology Shared Resource, the Microscopy Core, the Metabolomics Core, and the Translational Tissue Modeling Laboratory, all at the University of Michigan. We also thank Peter Kuffa, Yadong Mao, and Ingrid L. Bergin for experimental assistance and Eric C Martens for providing bacterial strains of SM. This work was supported by National Institutes of Health grants DK110146, DK108901, DK125087, DK119219, and AI142047 (to N.K.), the Kenneth Rainin Foundation Innovator Award and Synergy Award (to N.K.), a JSPS Postdoctoral Fellowship for Research Abroad (to K.S., S.K., and H.N.-K.), the Uehara Memorial Foundation Postdoctoral Fellowship Award (to K.S., and S.K.), and the Crohn’s and Colitis Foundation (to K.S., H.N.-K., and N.K.), and the Office of the Assistant Secretary of Defense for Health Affairs endorsed by the Department of Defense through the Peer-Reviewed Cancer Research Program under Award No. W81XWH2010547 (to S.K.).

## Author contributions

K.S and N.K. conceived and designed the experiments. K.S. conducted most of the experiments with help from S.K., P.S., H.N-K., A.R., J.I., and M.O.. M.G.G. and N.I. performed 16S rRNA sequencing and RNA sequencing analysis. P.S., C.L.M., C.K.M., M.H., A.N., and J.G. contributed to the establishment of the colonoid culture and anaerobic coculture experiment. S.B., J.Y.K., C.J.A., N.B., A.N., and J.G. provided advice and constructive discussion of the results. K.S. and N.K. analyzed the data. K.S. and N.K. wrote the manuscript with contributions from all authors.

## Competing interests

The authors declare no competing interests.

## Extended Data Figure Legends

**Extended Data Fig. 1. IBD-associated AIEC LF82 preferentially uses L-serine for its growth**.

AIEC strain LF82 was cultured in DMEM/F12 media for 1 or 3 h and then the concentrations of amino acids were measured. Dots indicate individual samples with mean ± SEM.. \**P* < 0.05, \*\**P* < 0.01, \*\*\**P* < 0.001 by 1-way ANOVA with Tukey post hoc test.

**Extended Data Fig. 2. Dietary L-serine restriction-induced exacerbation of colitis is dependent on the gut microbiota**.

**a**, SPF C57BL/6 mice were treated with drinking water containing a cocktail of antibiotics (ampicillin, neomycin, vancomycin) and metronidazole by oral gavage for 7 days. Mice were then fed the Ctrl diet or the ΔSer diet and treated with DSS for 5 days. During DSS treatment, mice were given a cocktail of antibiotics (ampicillin, neomycin, vancomycin, metronidazole) by oral gavage. On day 5 post-DSS, all mice were euthanized. **b**, Bacterial burden of feces after treatment with antibiotics was evaluated by qPCR. **c**, Body weight was measured during DSS treatment. **d–f**, colonic length, representative histological images of HE sections (scale bar, 200 μm), and histological scores. Dots indicate individual mice, with mean ± SEM. NS, not significant. \*\*\**P* < 0.001 by unpaired *t* test.

**Extended Data Fig. 3. AIEC and *A. muciniphila* enhance gut inflammation in a dietary L-serine–dependent manner**.

**a**, Experimental protocol and the composition of the nonmucolytic synthetic human gut microbiota (NmSM) for the gnotobiotic mouse experiments. **b,c**, AB/PAS staining and immunofluorescence staining of colonic sections (MUC2, green; DAPI, blue). The thickness of the inner mucus layer was measured (scale bar, 100 μm). **d**, Homogenates of colon tissues were cultured on LB agar plates. The number of viable bacteria was estimated by counting the CFUs. **e,f**, Body weight and colon length were measured 7 days post-DSS treatment. **g,h**, Representative images of colonic sections stained with HE (scale bar, 200 μm) and histology scores. **i**, Intestinal permeability was assessed with FITC–dextran. Dots indicate individual mice, with mean ± SEM. NS, not significant. \**P* < 0.05, \*\**P* < 0.01, \*\*\**P* < 0.001 by 1-way ANOVA with Tukey post hoc test.

**Extended Data Fig. 4. The transcriptomic profiles of AIEC and the human-derived colonoid monolayer (HCM)**.

**a**, Experimental design. LF82 was cultured with or without HCM for 3 h. Uninfected HCM was used as a control to assess the impact of LF82 infection in transcriptome of HCM. **b**, Volcano plot shows the significantly upregulated and downregulated genes in LF82-associated with HCM (fold change >2 and SNR >1). **c**, Significantly upregulated and downregulated pathways of LF82 transcriptomes identified by GO enrichment analysis. **d**, Volcano plot shows the significantly up- and downregulated genes in the infected-HCM (fold change >2 and SNR >1). **e**, Significantly upregulated and downregulated pathways of HCM transcriptomes identified by GO enrichment analysis.

## Notes

### Competing Interest Statement

The authors have declared no competing interest.

### Summary of Updates

A contributing author added.

