## Supplemental Fig. 1 for "Mucolytic bacteria license pathobionts to acquire host-derived nutrients during dietary nutrient restriction"

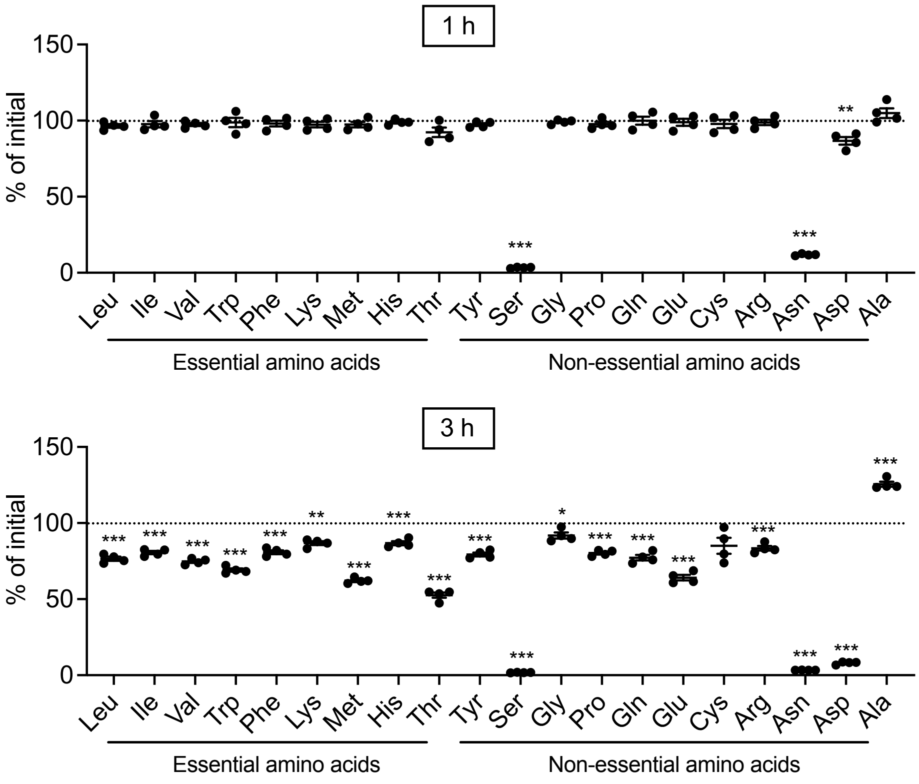


**Figure S1. IBD-associated AIEC LF82 preferentially uses L-serine for its growth, Related to Figure 1.**

AIEC strain LF82 was cultured in DMEM/F12 media for 1 or 3 h and then the concentrations of amino acids were measured. Dots indicate individual samples with mean ± SEM. . *p < 0.05, **p < 0.01, ***p < 0.001 by 1-way ANOVA with Tukey post hoc test.
