## Supplemental Fig. 2 for "Mucolytic bacteria license pathobionts to acquire host-derived nutrients during dietary nutrient restriction"

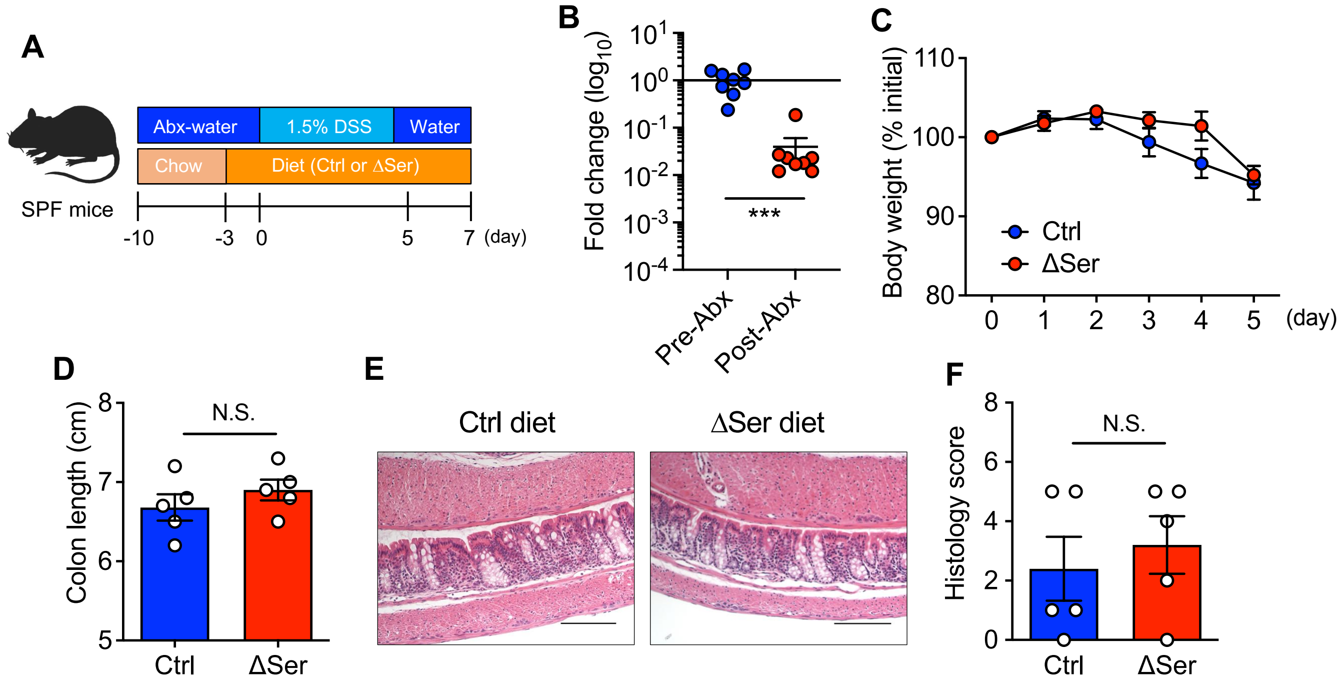


**Figure S2. Dietary L-serine restriction-induced exacerbation of colitis is dependent on the gut microbiota, Related to Figure 2**

(A) SPF C57BL/6 mice were treated with drinking water containing a cocktail of antibiotics (ampicillin, neomycin, vancomycin) and metronidazole by oral gavage for 7 days. Mice were then fed the Ctrl diet or the ΔSer diet and treated with DSS for 5 days. During DSS treatment, mice were given a cocktail of antibiotics (ampicillin, neomycin, vancomycin, metronidazole) by oral gavage. On day 5 post-DSS, all mice were euthanized.
(B) Bacterial burden of feces after treatment with antibiotics was evaluated by qPCR.
(C) Body weight was measured during DSS treatment.
(D–F) colonic length, representative histological images of HE sections (scale bar, 200 μm), and histological scores. Dots indicate individual mice, with mean ± SEM. N.S., not significant. ***p < 0.001 by unpaired *t* test.
