## Supplemental Fig. 3 for "Mucolytic bacteria license pathobionts to acquire host-derived nutrients during dietary nutrient restriction"

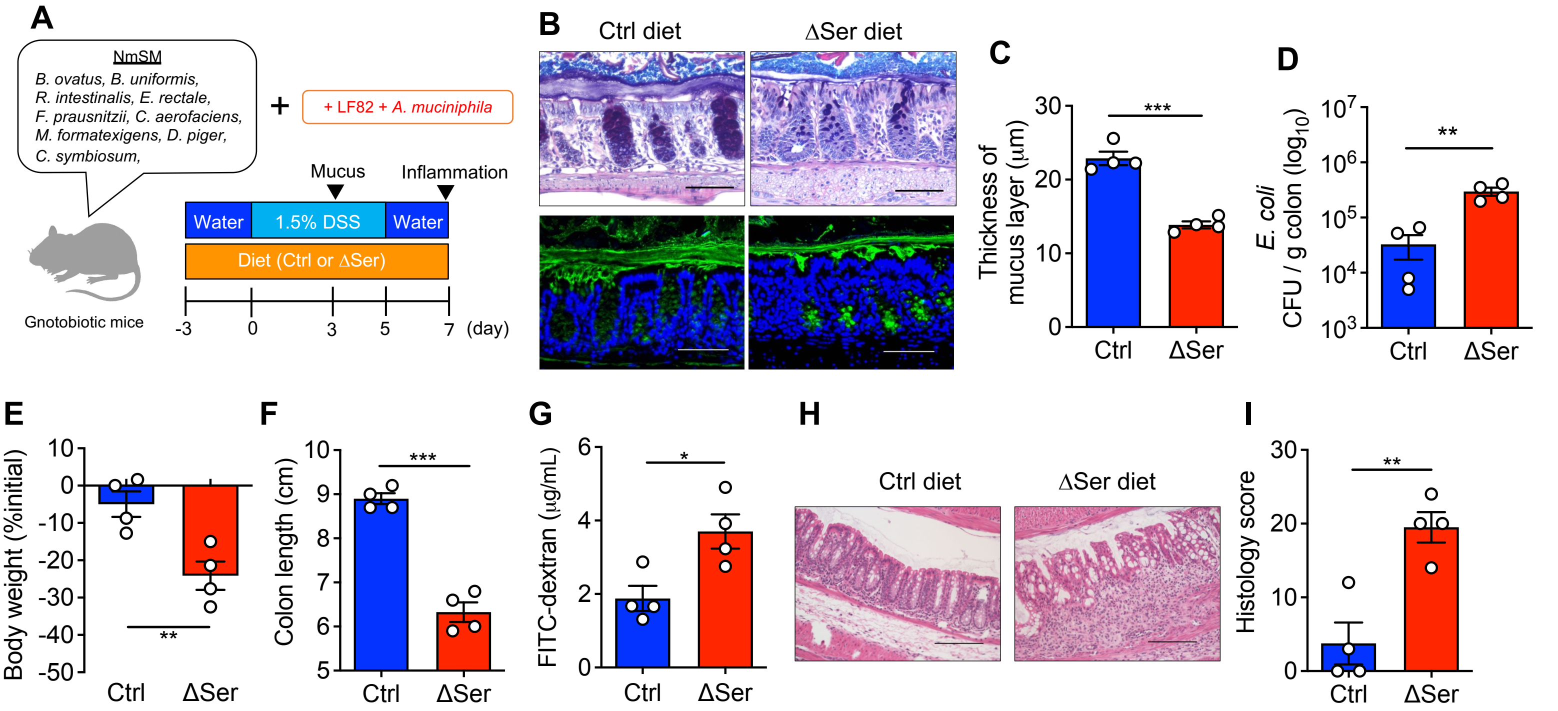


### Figure S3. AIEC and *A. muciniphila* enhance gut inflammation in a dietary L-serine–dependent manner, Related to Figure 6

(A) Experimental protocol and the composition of the nonmucolytic synthetic human gut microbiota (NmSM) for the gnotobiotic mouse experiments.
(B and C) AB/PAS staining and immunofluorescence staining of colonic sections (MUC2, green; DAPI, blue). The thickness of the inner mucus layer was measured (scale bar, 100 μm).
(D) Homogenates of colon tissues were cultured on LB agar plates. The number of viable bacteria was estimated by counting the CFUs.
(E and F) Body weight and colon length were measured 7 days post-DSS treatment.
(G and H) Representative images of colonic sections stained with HE (scale bar, 200 μm) and histology scores.
(I) Intestinal permeability was assessed with FITC–dextran.
Dots indicate individual mice, with mean ± SEM. N.S., not significant. *p < 0.05, **p < 0.01, ***p < 0.001 by 1-way ANOVA with Tukey post hoc test.
