## Supplemental Fig. 4 for "Mucolytic bacteria license pathobionts to acquire host-derived nutrients during dietary nutrient restriction"

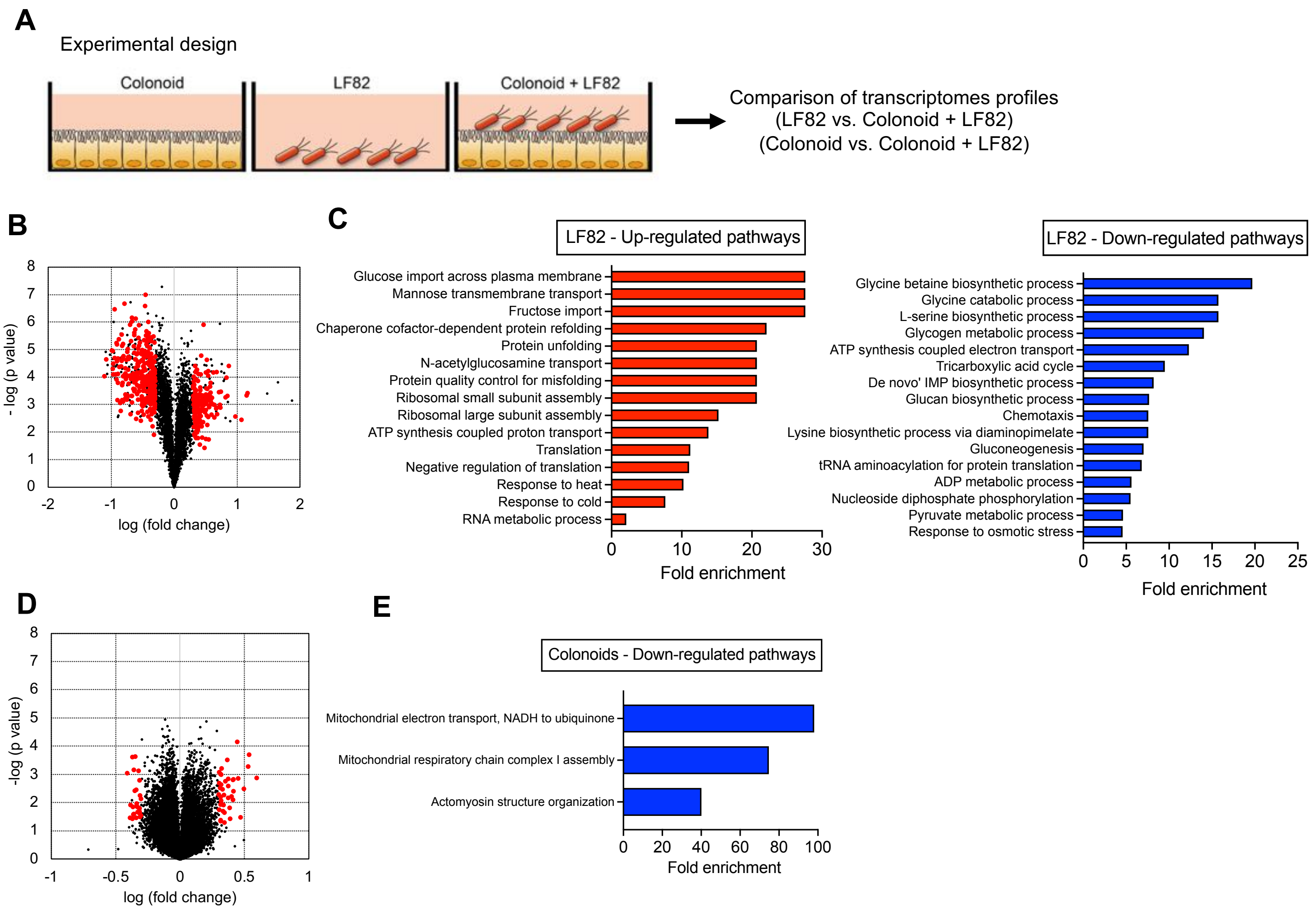


**Figure S4. The transcriptomic profiles of AIEC and the human-derived colonoid monolayer (HCM), Related to Figure 7**

(A) Experimental design. LF82 was cultured with or without HCM for 3 h. Uninfected HCM was used as a control to assess the impact of LF82 infection in transcriptome of HCM.
(B) Volcano plot shows the significantly upregulated and downregulated genes in LF82-associated with HCM (fold change >2 and SNR >1).
(C) Significantly upregulated and downregulated pathways of LF82 transcriptomes identified by GO enrichment analysis.
(D) Volcano plot shows the significantly up- and downregulated genes in the infected-HCM (fold change >2 and SNR >1).
(E) Significantly upregulated and downregulated pathways of HCM transcriptomes identified by GO enrichment analysis.
