## Supplemental Table 1 for "Mucolytic bacteria license pathobionts to acquire host-derived nutrients during dietary nutrient restriction"

**Supplementary Table 1: Primers used in this study.**

| Primer sequence | Source | Reference |
|---|---|---|
| Eubacteria16S F: 5'-ACTCCTACGGGAGGCAGCAGT-3' | Burr et al., 2006 | https://pubmed.ncbi.nlm.nih.gov/16645925/ |
| Eubacteria16S R: 5'-ATTACCGCGGCTGCTGGC-3' | | |
| B. ovatus F: 5'-GTGAAGGTGCCATCGGAGGAC-3' | Desai et al., 2016 | https://pubmed.ncbi.nlm.nih.gov/27863247/ |
| B. ovatus R: 5'-GGACGCTTTGGCCACTATTTCA-3' | | |
| B. uniformis F: 5'-GCTACCGGGAGATACTGGATTGG-3' | Desai et al., 2016 | https://pubmed.ncbi.nlm.nih.gov/27863247/ |
| B. uniformis R: 5'-TGCGGCGGCCTTTGAAC-3' | | |
| E. rectale F: 5'-AGCTTGTGCCGCCCATCTCTAT-3' | Desai et al., 2016 | https://pubmed.ncbi.nlm.nih.gov/27863247/ |
| E. rectale R: 5'-TTGCGGTAAAGCTTTGGTGTGG-3' | | |
| C. symbiosum F: 5'-CCGCTTGGCATGAAACAGGTATC-3' | Desai et al., 2016 | https://pubmed.ncbi.nlm.nih.gov/27863247/ |
| C. symbiosum R: 5'-TTGGAAGCGGCGAAGAATGG-3' | | |
| E. coli F: 5'-GGTGGCTGGGTGATGTAAAACTGA-3' | Desai et al., 2016 | https://pubmed.ncbi.nlm.nih.gov/27863247/ |
| E. coli R: 5'-ACCGCCGAGCAAAATGAAGC-3' | | |
| A. municiphila F: 5'-GACCGGCATGTTCAAGCAGACT-3' | Desai et al., 2016 | https://pubmed.ncbi.nlm.nih.gov/27863247/ |
| A. municiphila R: 5'-AAGCCGCATTGGGATTATTTGTT-3' | | |
| R. intestinalis F: 5'-TCGAAATTAAAGAGACGGAAACAGAAG-3' | Desai et al., 2016 | https://pubmed.ncbi.nlm.nih.gov/27863247/ |
| R. intestinalis R: 5'-CCGCTCATATCAATCGAAAACACA-3' | | |
| F. prausnitzii F: 5'-TGCCCCCGGGTGGTTCT-3' | Desai et al., 2016 | https://pubmed.ncbi.nlm.nih.gov/27863247/ |
| F. prausnitzii R: 5'-CGTTATTCAAAGCCCCGTTATCAA-3' | | |
| M. formatexigens F: 5'-CAGGGATTTTACGTGCTTTATTTTAGTTAT-3' | Desai et al., 2016 | https://pubmed.ncbi.nlm.nih.gov/27863247/ |
| M. formatexigens R: 5'-AGTTCGGATTCGCTCGTATTTTCT-3' | | |
| C. aerofaciens F: 5'-GTGCGGCCGAAAACCAAATG-3' | Desai et al., 2016 | https://pubmed.ncbi.nlm.nih.gov/27863247/ |
| C. aerofaciens R: 5'-CCACGCGCAGGAGCAAAAA-3' | | |
| D. piger F: 5'-TGGCTTCAGGCAAATCTCAAAT-3' | Desai et al., 2016 | https://pubmed.ncbi.nlm.nih.gov/27863247/ |
| D. piger R: 5'-TCCGGGGAATCAAAACCATAC-3' | | |
| E. coli_fimH F: 5'-TGCCGTGCTTATTTTGCGAC-3' | this study | |
| E. coli_fimH R: 5'-GGCACTGAACCAGGGTAGTC-3' | | |
| E. coli_ibeA F: 5'-AGAAACGGCAAAATCAATGG-3' | this study | |
| E. coli_ibeA R: 5'-TGATAACATCAACGGCGGTA-3' | | |
| E. coli_ompA F: 5'-TGGGTGTTTCCTACCGTTTC-3' | Hejair et al., 2017 | https://pubmed.ncbi.nlm.nih.gov/28315387/ |
| E. coli_ompA R: 5'-GAGTGAAGTGCTTGGTCTGT-3' | | |
| E. coli_ompC F: 5'-TTCTTCGGTCTGGTTGACGG-3' | this study | |
| E. coli_ompC R: 5'-ATTTTCACCGCTTACGCTGC-3' | | |

| | | |
|---|---|---|
| E. coli_vat F: 5’-ATAGCATCCTGCCCTCTTTTGG-3’ | Gibold et al., 2016 | https://pubmed.ncbi.nlm.nih.gov/26499863/ |
| E. coli_vat R: 5’-CTGTGAGAGAAAACTCTGAGG+C57-3’ | | |
| E. coli_fliC F: 5’-TGGTGCTGCAACTGCTAACGC-3’ | Barnich et al., 2004 | https://pubmed.ncbi.nlm.nih.gov/15102755/ |
| E. coli_fliC R: 5’-TTATCGGCATATTTTGCGCTAGC-3’ | | |
| E. coli_pduC F: 5’-CCTGCAAAAATCGTCGAAGTGGTT-3’ | Dogan et al., 2014 | https://pubmed.ncbi.nlm.nih.gov/25230163/ |
| E. coli_pduC R: 5’-TTGGTGACGTGAGCCTGTTGTGAT-3’ | | |
| E. coli_chuA F: 5’-ATGGTACCGGACGAACCAAC-3’ | Hashimoto et al., 2021 | https://pubmed.ncbi.nlm.nih.gov/33540892/ |
| E. coli_chuA R: 5’-TGCCGCCAGTACCAAAGACA-3’ | | |
| E. coli_fyuA F: 5’-GGCGGCGTGCGCTTCTCGCA-3’ | Dezfulian et al., 2003 | https://pubmed.ncbi.nlm.nih.gov/12682117/ |
| E. coli_fyuA R: 5’-CGCAGTAGGCACGATGTTGTA-3’ | | |
